# AT7867 promotes pancreatic progenitor differentiation of human iPSCs and accelerates diabetes reversal

**DOI:** 10.1101/2023.05.11.538780

**Authors:** Nerea Cuesta-Gomez, Kevin Verhoeff, Nidheesh Dadheech, Rena Pawlick, Braulio Marfil-Garza, Haide Razavy, A.M. James Shapiro

## Abstract

Generation of pure pancreatic progenitor cells (PPs) is critical for clinical translation of stem cell derived islets. Herein, we performed PP differentiation with and without AKT/P70 inhibitor AT7867 and characterized the resulting cells at protein and transcript level *in vitro* and *in vivo* upon transplantation into diabetic mice. AT7867 treatment increased the percentage of PDX1^+^NKX6.1^+^ (-AT7867: 50.9% [IQR 48.9%-53.8%]; +AT7867: 90.8% [IQR 88.9%-93.7%]; *p=0.0021*) and PDX1^+^GP2^+^ PP cells (-AT7867: 39.22% [IQR 36.7%-44.1%; +AT7867: 90.0% [IQR 88.2%-93.6%]; *p=0.0021*). Transcriptionally, AT7867 treatment significantly upregulated *PDX1* (*p=0.0001*), *NKX6.1* (*p=0.0005*) and *GP2* (*p=0.002*) expression compared to controls, while off-target markers *PODXL* (*p<0.0001*) and *TBX2* (*p <0.0001*) were significantly downregulated. Transplantation of AT7867 treated PPs resulted in faster hyperglycemia reversal in diabetic mice compared to controls (time and group: *p<0.0001*). Overall, our data shows that AT7867 enhances PP cell differentiation leading to accelerated diabetes reversal.

## 1 Introduction

Islet cell transplantation has demonstrated proof-of-concept that β-cell replacement therapies have the potential to improve glycemic control in patients with diabetes ^1^. Limited donor supply remains a major limitation to widespread islet transplantation and pluripotent stem cells offer a renewable source for the generation of stem cell-derived islets (SC-islets) ^2–5^. Through the addition of growth factors and small molecules, pluripotent stem cells can recapitulate embryological development and generate pancreatic progenitors (PP), which can further differentiate *in vivo* and reverse diabetes ^6^. Alternatively, PPs can be further differentiated down the β-cell pathway and generate SC-islets *in vitro* ^7^. However, due to the trilineage differentiation potential of pluripotent stem cells, a major challenge hampering clinical translation of SC-islets is the heterogeneity of the cells generated, resulting in proliferative non-endocrine cell populations ^8–12^. Protocols to improve efficiency of PP or SC-islet differentiation are required to eliminate off-target populations and enable clinical translation.

To improve SC-islet differentiation and avoid non-pancreatic populations, endodermal commitment defined by CD184, CD117 and SOX17 expression^13,14^, as well as early and rigorous patterning of cells for pancreatic linage commitment (FOXA2^+^PDX1^+^) is critical^15–17^. PP generation requires the co-expression of key transcription factors, including PDX1, NKX6.1 and GP2^13,18,19^. Additionally, expression of Neurogenin-3 (NEUROG3)^20^ followed by the expression of mature endocrine markers such as chromogranin A (CHGA), NEUROD1 and NKX2.2 leads to progenitors committed to the endocrine lineage^21–24^. Alternatively, cell sorting, disaggregation and re-aggregation, and gene editing have been explored to remove off-target cell populations following non-specific differentiation^13,16,25,26^. Unfortunately, these approaches result in substantial cell loss and/or are not economically feasible^27^. Thus, we believe that optimization of the differentiation protocol to ensure only activation of pathways resulting in pancreatic and endocrine commitment is essential and we propose that at the PP stage, stage 4, at least 90% of the cells should express PDX1, NKX6.1 and GP2. Several small molecules, including nicotinamide^28,29^, TPB^29,30^ or Sant-1^9,31,32^ have previously been described to improve the generation of PDX1^+^NKX6.1^+^ PP cells. Similarly, preliminary studies have suggested AKT-inhibitor AT7867 may induce proliferation of PDX1^+^NKX6.1^+^ cells but the impact on PP maturation or *in vivo* PP maturation was not evaluated^33^. In addition, active AKT in PDX1^+^ PPs induces the proliferation of ductal structures, resulting in malignant lesions^34^. Hence, Thorough evaluation of the effect of AT7867 on PP proliferation, differentiation, and ensuing *in vivo* maturation is required to evaluate its potential to optimize PP generation.

This study aims to characterize the PP cells generated through the addition of AT7867 to a previously published differentiation protocol at transcript and protein level as well as to evaluate the potential of AT7867 treated PP cells to mature *in vivo* and reverse diabetes in mice.

## 2 Results

### 2.1 AT7867 increases the percentage of PDX1^+^NKX6.1^+^ and PDX1^+^GP2^+^ cells

We utilized a previously published protocol^35^ modified with addition of AT7867 to differentiate iPSCs into PP cells and evaluated the impact of AT7867 on PP cell composition and heterogeneity at stage 4 (**Figure 1A**). Prior to differentiation, iPSCs had compact cell-to-cell connections and condensed nucleus with minimum cytoplasm and 99.1% (IQR 98.8%-99.2%) and 98.9% (IQR 98.6%-99.4%) of the iPSCs were Oct4^+^SSEA4^+^ and Sox2^+^Nanog^+^, respectively (**Figure 1B**). Upon differentiation of iPSCs into definitive endoderm cells, morphologically, cells showed cytoplasmic enlargement and cell spacing compared to iPSCs; at this stage, 99.1% (IQR 98.8%-99.2%), 97.5% (IQR 96.9%-97.9%) and 96.6% (IQR 95.8%-97.2%) of the cells were CD184^+^CD117^+^, CD117^+^SOX17^+^ and CD184^+^CD117^+^Sox17^+^, respectively (**Figure 1C**). Further differentiation into primitive gut tube resulted in elongation of the cells and induction of FoxA2, where 65.9% (IQR 64.7%-68.5%) of cells were Sox17^+^FoxA2^+^(**Figure 1D**). Differentiation into posterior foregut resulted into further elongation of the cells with enlarged cytoplasm coupled with PDX1 induction, resulting in 86.6% (IQR 83.5%-88.1%) PDX1^+^ cells and 64.6% (IQR 63.5%-65.8%) FOXA2^+^PDX1^+^ cells. At stage 4, addition of AT7867 resulted in the formation of a homogeneous cell layer while the control cells (-AT7867) showed a monolayer of cells with breaks in the monolayer as a result of the formation of raised “ribbons” of cells (**Figure 1E**). Quality control of the PP cells using flow cytometry showed that 50.9% (IQR 48.9%-53.8%) and 39.2% (IQR 36.7%-44.1%) of the control -AT7867 PP were PDX1^+^NKX6.1^+^ and PDX1^+^GP2^+^ compared to 90.8% (IQR 88.9%-93.7%; *p=0.0021*) and 90.0% (IQR (88.2%-93.6%; *p=0.0021*) in AT7867 treated PPs, respectively. Furthermore, AT7867 treatment resulted in a reduced percentage of NKX6.1^+^CHGA^+^ cells (-AT7867: 6.0% [IQR 4.5%-8.4%]; +AT7867: 2.7% [IQR 2.4%-4.4%]; *p=0.0151*). Increased PDX1^+^NKX6.1^+^ and PDX1^+^GP2^+^ upon addition of AT7867 has been tested in three independent iPSC lines showing similar results (**Figure S1**).

**Figure 1.**
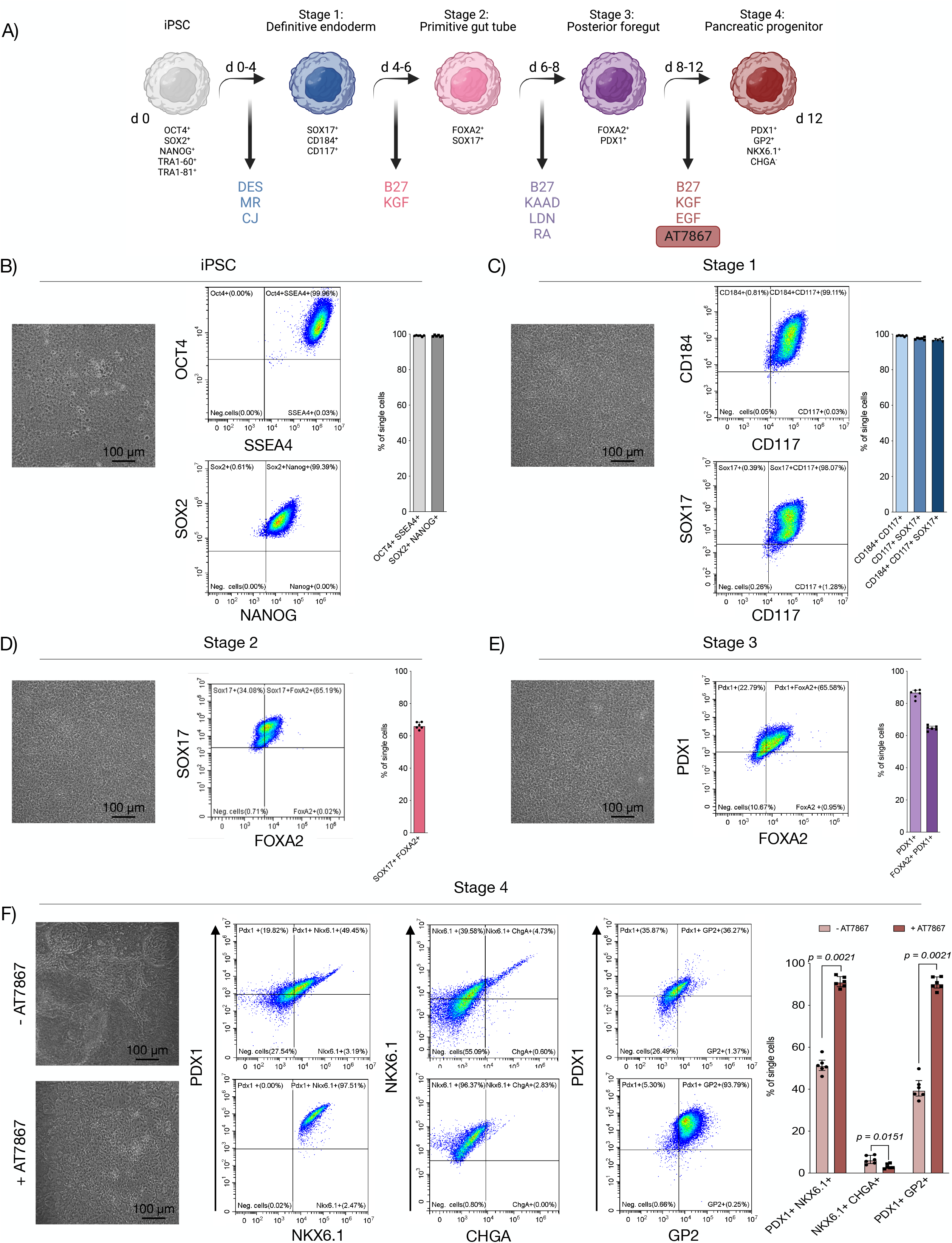
AT7867 increases the percentage of Pdx1^+^Nkx6.1^+^ and Pdx1^+^GP2^+^ cells. A) Schematic representation of the differentiation protocol. B) Microscopy of iPSC culture and flow cytometry analysis of iPSCs with quantification. C) Representative microscopy and flow cytometric analysis with quantification of cells at stage 1, D) stage 2, E) stage 3 and F) stage 4 of 6 independent experiments. All data are represented as median with IQR.

### 2.2 AT7867-mediated PDX1^+^NKX6.1^+^ cell population increase is not a result of pancreatic progenitor proliferation

In order to assess the mechanism by which AT7867 increased PDX1 and NKX6.1 expression we evaluated whether selective proliferation of PP cells occurred upon addition of AT7867. First, we quantified the number of cells at the end of the PP stage of differentiation in control and treated cells and observed no differences (-AT7867: 8.7×10^6^ cells [IQR 8.1×10^6^-9.0×10^6^ cells]; +AT7867: 8.5×10^6^ cells [IQR 8.3×10^6^-9.3×10^6^ cells]; *p=0.6991*) (**Figure 2A**). Next, we performed western blotting to quantify the relative expression of PDX1 and KI67 present in 30 μg of protein; iPSCs were used as positive control for proliferation (Ki67) and as negative control for PDX1 expression (**Figure 2B**). No differences were observed in the relative density of KI67 between control and AT7867 treated PP cells (-AT7867: 1.07 [IQR 1.02-1.1]; +AT7867: 1.07 [IQR 1.05-1.2]; *p>0.9999*) (**Figure 2C**). However, the relative density of PDX1 was significantly increased in AT7867 treated PP cells compared to control (-AT7867: 3130 [IQR 1528-3294]; +AT7867: 5975 [IQR 4667-6511]; *p=0.0375*) (**Figure 2D**). These results were further confirmed with immunohistochemistry (**Figure 2E** and **S2A**), where no differences were observed upon the quantification of positive cells stained for KI67 (-AT7867: 80.9% [IQR 66.7%-92.4%]; +AT7867: 91.0% [IQR 70.9%-95.4%]; *p=0.5054*), despite statistically more PDX1^+^ cells upon treatment with AT7867 (-AT7867: 43.6% [IQR 38.5%-50.0%]; +AT7867: 79.9% [IQR 69.1%-85.9%]; *p=0.0002*) (**Figure 2F**). Furthermore, no differences were observed in the percentage of KI67+ cells within the PDX1+ population between control and AT7867 treated PPs (-AT7867: 99.55% [IQR 99.07%-99.98%]; +AT7867: 99.16% [IQR98.89%-99.60%]; *p=0.3283*; **Figure 2G**). Importantly, the percentage of KI67^+^ cells using flow cytometry daily throughout stage 4 (**Figure 2H** and **S2B**), showed no differences in proliferation between control versus AT7867 treated cells (medians and IQR can be found in **Table S1)**.

**Figure 2.**
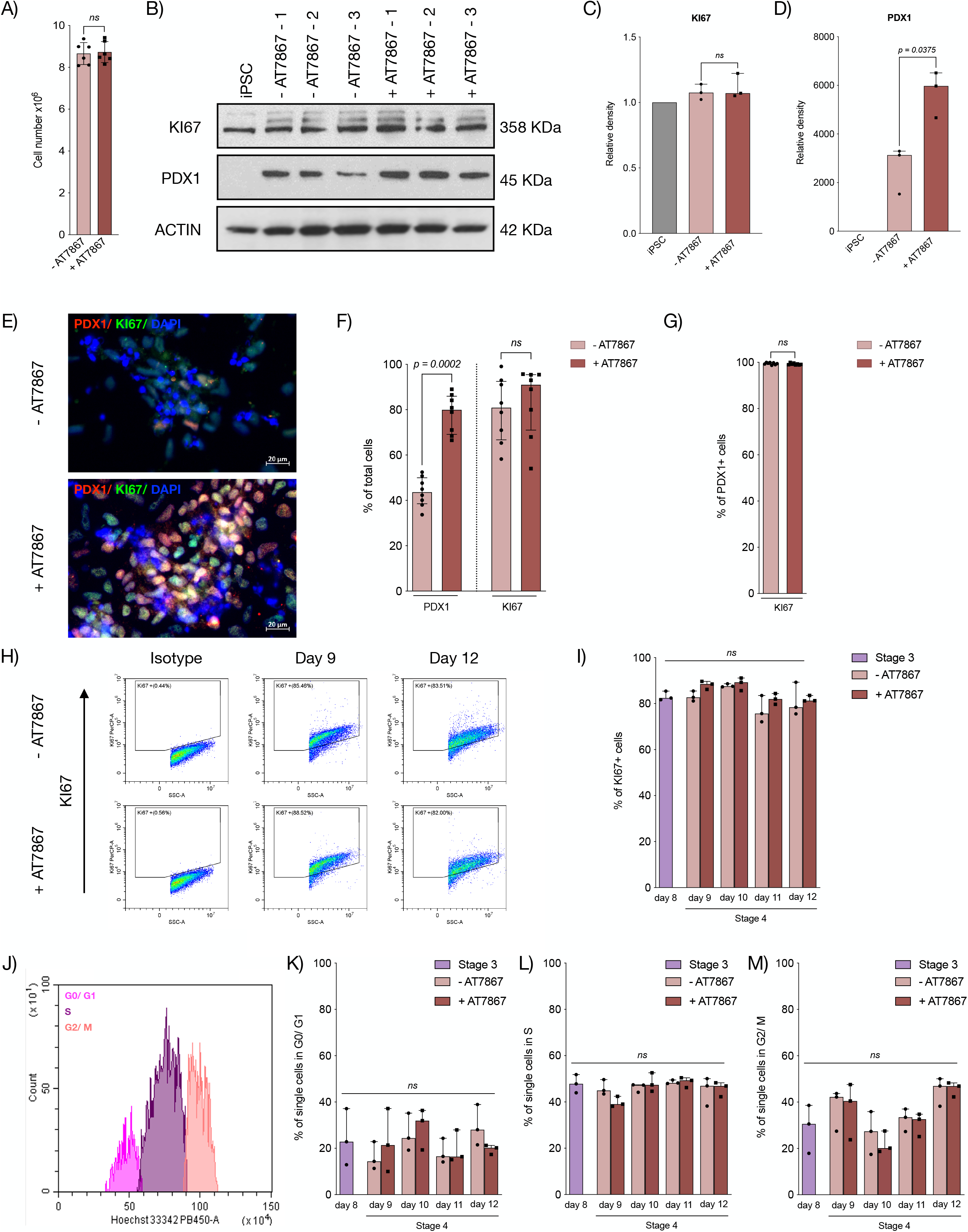
AT7867 does not induce the proliferation of pancreatic progenitor cells. A) Quantification of the cell number at the end of stage 4. B) Representative western blot analyzing KI67 and PDX1 protein content. C) Quantification of the relative density of Ki67 and D) PDX1 from three independent experiments. E) Representative immunohistochemistry of PDX1 and KI67 expression. F) Quantification of the percentage of total cells positive for PDX1 and KI67 from 8 independent experiments. G) Quantification of the percentage of PDX1^+^ and PDX1^-^ cells within the KI67^+^ population from 8 independent experiments. H) Representative flow cytometry analysis and quantification of KI67^+^ cells. I) Percentage of single KI67^+^ cells from 3 independent experiments. J) Representative flow cytometry gating of cells in G0/G1, S and G2/M phases. K) Percentage of single cells in G0/G1, L) S and M) G2/M phase throughout stage 4 from 3 independent experiments. All data are represented as median with IQR.

Finally, daily analysis of the cell cycle using DNA staining with Hoescht 33342 dye demonstrated a clear delineation of cells in the G0/G1 phase, S phase, and G2 and M phases to further quantify proliferation. (**Figure 2J**). No statistically significant differences were observed in the percentage of cells undergoing G0/G1 phase (**Figure 2K**), S phase (**Figure 2L**) or G2/M phase (**Figure 2M**) throughout stage 4 between AT7867 treated PP cells and control PP cells; medians and IQR can be found in **Table S2**).

In summary, while AT7867 enriches the PDX1^+^NKX6.1^+^GP2^+^ population, it appears that these differences are not a result of AT7867-mediated increased cell proliferation evaluated by KI67, or cell cycle quantification.

### 2.3 AT7867 induces the transcriptional upregulation of genes associated with pancreatic progenitor and pancreatic endocrine lineage commitment

Our next step to evaluate the mechanism by which AT7867 increased the number of PDX1^+^NKX6.1^+^GP2^+^ cells was to analyze the expression of genes associated with PP development and pancreatic endocrine progenitor (PEP) lineage commitment.

Transcriptome analysis represented as a heatmap showcased key differences in the transcription of key genes involved in PP differentiation and PEP lineage specification (**Figure 3A).** Specifically, genes associated with pancreatic progenitor commitment (*FOXA2, GP2, NKX6.1, ONECUT1* and *PDX1*) were upregulated in AT7867 treated PP cells compared to control PP cells (**Figure 3A** and **S3**). Similarly, genes associated with PEP lineage commitment (*ARX, CHGA, HNF4A, ISL1, MAFB, NEUROD1, NEUROG3, NKX2.2, PAX6* and *UCN3*) and genes associated with endocrine cell maturation or hormone secretion (*GCG, INS, PAX4, PCSK1* and *SST*) were upregulated in AT7867 treated PP cells compared to control PP cells (**Figure 3A** and **S3**). On the contrary, the expression of β-cell identity markers *SLC30A8, PCSK2* and *TSPAN1* was downregulated in AT7867 treated PP compared to control PP cells (**Figure 3A** and **S3**). Hierarchical clustering using the complete clustering method showed similarities between biological replicates collected for each condition, validating the reproducibility of the expression data.

**Figure 3.**
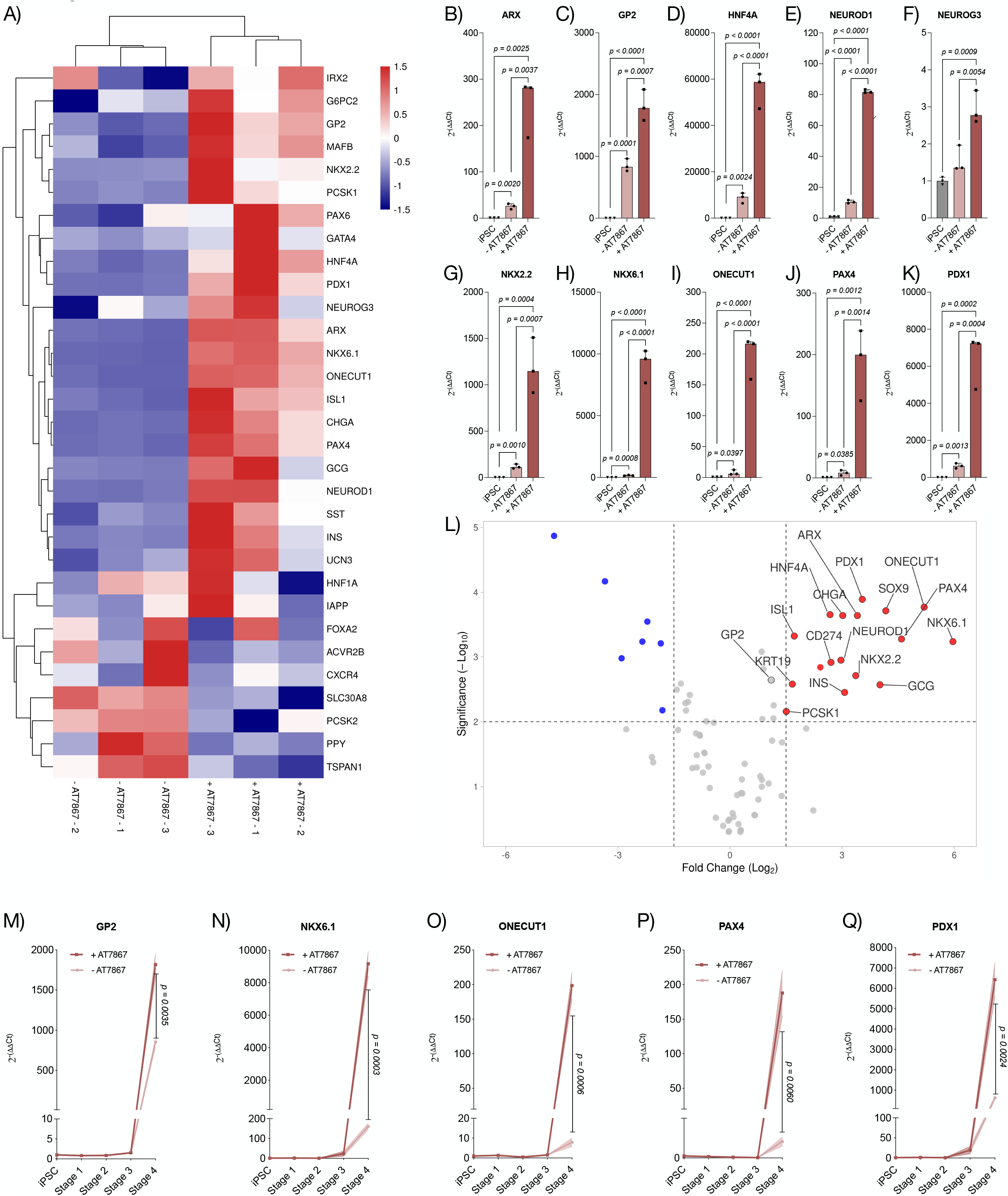
AT7867 induces the transcriptional upregulation of genes associated with pancreatic progenitor and pancreatic endocrine commitment. A) Heatmap representation of transcript levels from control and AT7867 treated PPs. B) Representation of 2^-ΔΔCt^ of *ARX*, C) *GP2,* D) *HNF4A*, E) *NEUROD1*, F) *NEUROG3*, G) *NKX2.2*, H) *NKX6.1*, I) *ONECUT1*, J) *PAX6* and K) *PDX1* in control and AT7867 treated PPs. L) Volcano plot representation of the transcriptome of AT7867 treated PPs vs. controls. M) Representation of the transition of the expression of *GP2*, N) *NKX6.1*, O) *ONECUT1*, P) *PAX4* and Q) *PDX1* from undifferentiated iPSC to PP for control and AT7867 treated PPs. All data are represented as median with IQR from 3 independent experiments.

Analysis of the fold change represented as 2^-(ΔΔCt)^ showed that the above-mentioned genes were all upregulated in both AT7867 treated and control PPs compared to undifferentiated iPSCs. However, addition of AT7867 resulted in a statistically significant upregulation of PP commitment genes (*GP2* [**Figure 3C**], *NKX6.1* [**Figure 3H**]*, ONECUT1* [**Figure 3I**] and *PDX1* [**Figure 3K**]), PEP commitment genes *(ARX* [**Figure 3B**]*, HNF4A* [**Figure 3D**]*, NEUROD1* [**Figure 3E**]*, NEUROG3* [**Figure 3F**]*, NKX2.2* [**Figure 3G**] and endocrine cell maturation marker *PAX4* [**Figure 3J**]) compared to controls. Median, IQR and statistical significance for these genes can be found in **Table S3**.

Volcano plot visualization of genes with statistically significant fold changes in AT7867 treated PP cells compared to control PP cells further confirmed that the expression of genes associated with PP commitment (*GP2, NKX6.1, ONECUT1* and *PDX1*), PEP lineage (*ARX, CHGA, HNF4A, INS, ISL1, NEUROD1, NEUROG3, NKX2.2, NKX6.1, ONECUT1* and *PDX1*) and hormone secretion (*GCG, INS, PAX4* and *PCSK1*) were significantly upregulated (*p<0.01*) in AT7867 treated PP cells compared to control PP cells (**Figure 3L**). Fold change and p value of all the genes represented in the Volcano plot can be found in **Table S4**.

Lastly, analysis of the expression of *GP2* (**Figure 3M**), *NKX6.1* (**Figure 3N**), *ONECUT1* (**Figure 3O**), *PAX4* (**Figure 3P**) and *PDX1* (**Figure 3Q**) throughout differentiation (iPSC to PP) in control and AT7867 treated samples showed a sharp upregulation of these transcripts at stage 4; furthermore, this upregulation throughout differentiation was significantly increased upon treatment with AT7867 at stage 4 (*GP2*: *p=0.0035*; *NKX6.1*: *p=0.0003*; *ONECUT1*: *p=0.0006*; P*AX4*: *p=0.0060* and *PDX1*: *p=0.0024*).

In summary, addition of AT7867 at stage 4 resulted in the upregulation of key genes involved in PP and PEP lineage commitment as well as hormone secretion. AT7867 therefore improves PDX1^+^NKX6.1^+^ phenotype acquisition by directly improving differentiation efficiency.

### 2.4 AT7867 induces the transcriptional downregulation of genes associated with pluripotency and non-endocrine populations

To characterize the impact of improved differentiation with AT7867 on non-endocrine cell populations we compared AT7867 treated PPs to control PPs. To accomplish this, we evaluated the expression of genes associated with pluripotency, early stages of differentiation or non-pancreatic endocrine populations.

Transcriptome analysis of key genes involved in the establishment and maintenance of pluripotency as well as non-pancreatic endoderm populations showcased key differences in the transcription of these genes upon treatment with AT7867 (**Figure 4A).** Specifically, pancreatic ductal lineage markers *KRT19* and *SOX9*, enterochromaffin cell identity gene *SLC18A1* and neuroendoderm marker *GDF3* were upregulated in AT7867 treated PP cells compared to controls. On the other hand, expression of genes associated with pluripotency, including *MYC, KIT, PODXL, LIN28A* and *TERT,* as well as the mesenchymal marker *TBX2* were downregulated in AT7867 treated PP compared to control PP cells (**Figure 4A**, **S3** and **S4**). No statistically significant differences were observed in the expression of pluripotency markers *KLF4, ABCG2, PODXL2, SOX2, FUT4, CDH1, POU5F1, TPBG, NANOG, UTF1* and *ZFP42* upon treatment with AT7867 (**Figure 4A** and **S4**). Hierarchical clustering using the complete clustering method showed similarities between biological replicates collected for each condition, validating the reproducibility of the expression data (**Figure 4A**).

**Figure 4.**
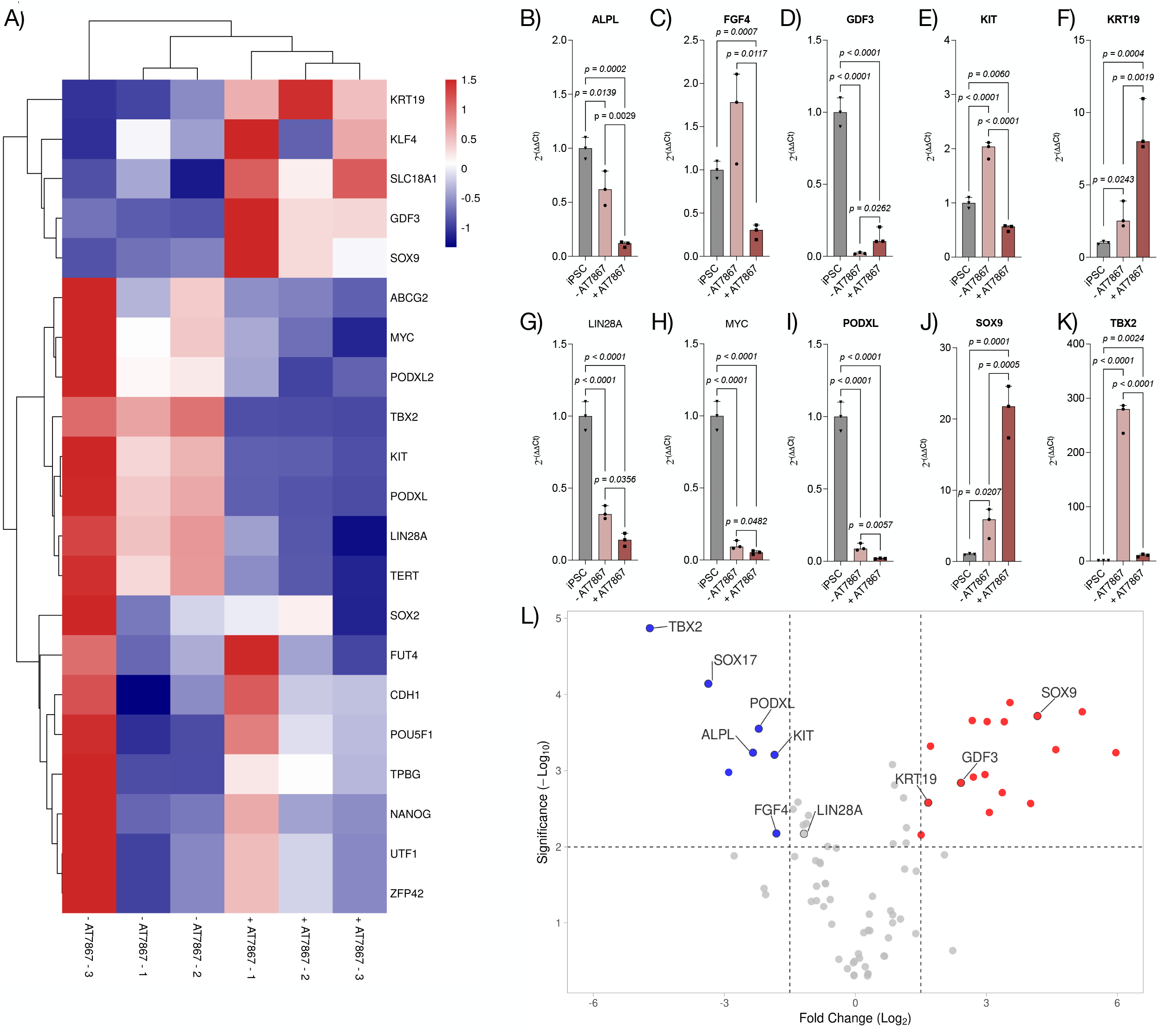
AT7867 induces the transcriptional downregulation of genes associated with pluripotency establishment and maintenance. A) Heatmap representation of transcript levels from control and AT7867 treated PPs. B) Representation of 2^-ΔΔCt^ of *ALPL*, C) *FGF4,* D) *GDF3*, *KIT*, F) *KRT19*, G) *LIN28A*, H) *MYC*, I) *PODXL*, J) *SOX9* and K) *TBX2* in control and AT7867 treated PPs. L) Volcano plot representation of the transcriptome of AT7867 treated PPs vs. controls. Data are represented as median with IQR from 3 independent experiments.

In addition, analysis of the fold change represented as 2^-(ΔΔCt)^ showed statistically significant downregulation of pluripotency genes *ALPL* (**Figure 4B**), *FGF4* (**Figure 4C**), *KIT* (**Figure 4E**)*, LIN28A* (**Figure 4G**)*, MYC* (**Figure 4H**) and *PODXL* (**Figure 4I**) as well as downregulation of the mesenchymal marker *TBX2* (**Figure 4K**) in AT7867 treated PPs compared to control PPs. On the contrary, pancreatic ductal lineage markers *KRT19* (**Figure 4F**) and *SOX9* (**Figure 4J**) were upregulated in AT7867 treated PPs compared to control PPs. Median, IQR and statistical significance for these genes can be found in **Table S3**.

Volcano plot visualization of genes with large fold changes that were statistically significant in AT7867 treated PP cells compared to control PP cells further confirmed downregulation of pluripotency markers *PODXL, ALPL, KIT, FGF4* and *LIN28A,* and mesenchymal marker *TBX2* (*p<0.01*). Expression of pancreatic ductal lineage markers *KRT19* and *SOX9* and neuroendoderm marker *GDF3* were significantly upregulated (*p<0.01*) (**Figure 4L**). Fold change and p value of all the genes represented in the Volcano plot can be found in **Table S4**.

In conclusion, addition of AT7867 at stage 4 resulted in the downregulation of key pluripotency genes. Furthermore, AT7867 treatment induced upregulation of genes associated with pancreatic ductal lineage, which also originates from PPs.

### 2.5 AT7867 treatment accelerates *in vivo* endocrine differentiation and diabetes reversal

To assess the potential of AT7867 treated PPs to undergo *in vivo* differentiation and reverse diabetes we expanded and differentiated iPSCs within clusters, with or without AT7867, using 0.1 L Vertical Wheel® bioreactors (VWB). The resulting PP clusters were transplanted under the kidney capsule of SCID beige immunodeficient diabetic mice (**Figure 5A**).

**Figure 5.**
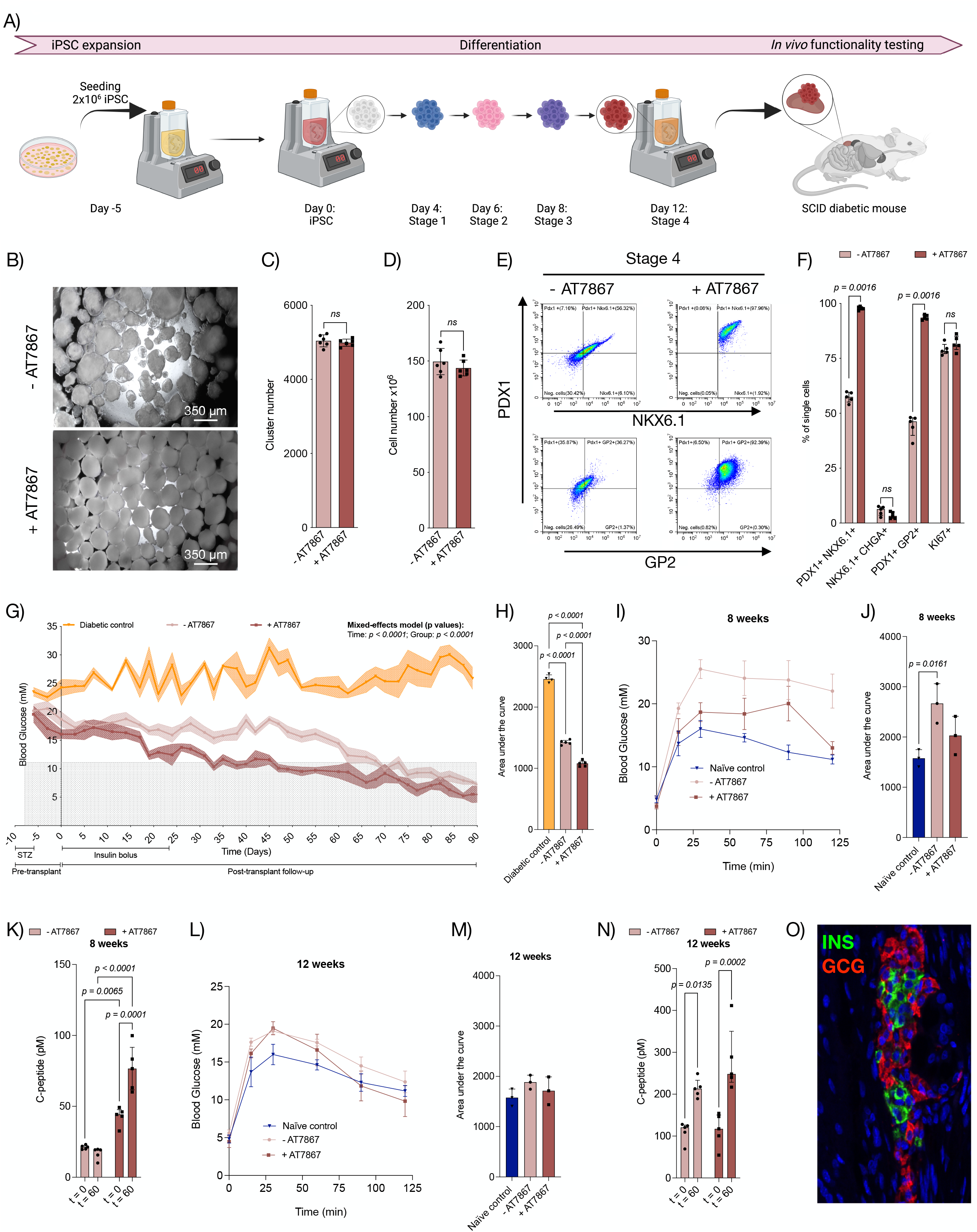
AT7867 treatment accelerates in vivo endocrine differentiation and diabetes reversal. A) Schematic representation of iPSC expansion and differentiation in 0.1 L VWB and *in vivo* functionality testing in SCID diabetic mice. B) Representative microscopy images of control and AT7867 treated PPs. C) Quantification of total cluster and D) total cell number at stage 4 from 6 independent differentiation experiments. E) Representative flow cytometry analysis and F) quantification of stage 4 markers from 6 independent experiments. G) Blood glucose measurements throughout experiment; 5 animals per group. H) Area under the curve measurements of the blood glucose readings per group (5 mice per group). I) Variations in glucose levels during IPGTT at 8 weeks. (J) Area under curve for IPGTT at 8 weeks. K) C-peptide concentration at t=0 and t=60 after glucose administration at 8 weeks. L) Variations in glucose levels during IPGTT at 12 weeks. (M) Area under curve for IPGTT at 12 weeks. N) C-peptide concentration at t=0 and t=60 after glucose administration at 12 weeks. O) Representative immunohistochemistry of the graft from mice transplanted with AT7867 treated PPs. All data are represented as median with IQR.

At the end of stage 4 differentiation, AT7867 treated PP clusters presented as tight clusters with a uniform and consistent aggregate size (278.8 μm [IQR 247.0-307.5 μm]) while control PP were bigger and more heterogenous in size (383.5 μm [IQR 293.8-459.7 μm]; *p<0.0001*) and morphology (**Figure 5B** and **S5**). No significant differences were observed in the number of clusters (-AT7867: 5038 [IQR 4915-5197]; +AT7867: 5002 [IQR 4904-5063]; *p=0.5887;* **Figure 5C**) or the number of cells (-AT7867: 147.0×10^6^ [IQR 139.4×10^6^-161.5×10^6^]; +AT7867: 141.4×10^6^ [IQR 137.9×10^6^-151.1×10^6^]; *p=0.5887;* **Figure 5D**) at the end of stage 4 differentiation with or without AT7867. Similar to cells differentiated on plates, quality control of the PP cells using flow cytometry showed that 57.4% (IQR 55.1%-59.5%) and 46.3% (IQR 39.9%-47.8%) of the control PP cells were PDX1^+^NKX6.1^+^ and PDX1^+^GP2^+^ compared to 97.6% (IQR 97.1%-98.5%; *p=0.0016*) and 93.5% (IQR (92.6%-94.7%; *p=0.0016*) in AT7867 treated PPs, respectively (**Figure 5E** and **5F**). Furthermore, AT7867 treatment had no effect on the number of NKX6.1^+^CHGA^+^ cells (-AT7867: 6.3% [IQR 3.5%-7.3%]; +AT7867: 3.2% [IQR 2.2%-5.0%]; *p=0.2030*) or proliferative KI67^+^ cells (-AT7867: 6.3% [IQR 3.5%-7.3%]; +AT7867: 3.2% [IQR 2.2%-5.0%]; *p=0.2244;* **Figure 5F**).

Glucose levels of the transplanted mice were monitored three times a week for 90 days to assess the potential of PP clusters to engraft, differentiate and reverse diabetes *in vivo* (**Figure 5G**). The diabetic control mice exhibited elevated glucose levels over 20 mM for the duration of the experiment. Mice transplanted with control AT7867 untreated PP clusters displayed gradual reversal of hyperglycemia over 70 days (IQR 66-78 days) and the animals remained non-diabetic. Interestingly, mice transplanted with AT7867 treated PP clusters displayed rapid reversal of hyperglycemia within 45 days (IQR 39-49 days; *p<0.0001*) of transplant and the animals remained non-diabetic over the remainder of the experiment. AT7867 treated PP clusters reversed diabetes significantly faster than controls (*p<0.0001*). This result was further confirmed by measurement of area under the curve (AUC) (-AT7867: 1423 [IQR 1390-1458]; +AT7867: 1080 [IQR 1033-1118]; *p<0.0001*; **Figure 5H**).

Intraperitoneal glucose tolerance test (IPGTT) performed at 8 weeks post-transplantation showed that, mice transplanted with AT7867 treated PP cells had IPGTT profiles more similar to naïve mice than mice transplanted with control PP clusters (**Figure 5I**). AUC of mice transplanted with AT7867 treated PP clusters was similar to naïve controls (Naïve control: 1577 [IQR 1406-1748]; +AT7867: 2029 [IQR 1648-2411]; *p=0.2897*) while mice transplanted with control PP clusters had a higher AUC (-AT7867: 2268 [IQR 2275-3061]; *p=0.0161;* **Figure 5J**). Mice transplanted with AT7867 treated PP clusters had increased concentration of C-peptide at time 0 compared to animals transplanted with control PP clusters (-AT7867 (t=0): 21.3 pM [IQR 19.7-22.4 pM]; +AT7867 (t=0): 44.29 pM [IQR 37.8-49.2 pM]; *p=0.0065*) (**Figure 5K**) and demonstrated glucose responsive C-peptide production 60 minutes after glucose administration (+AT7867 (t=0): 44.2 pM [IQR 37.8-49.2 pM]; +AT7867 (t=60): 76.6 pM [IQR 61.6-91.6 pM; *p=0.0001*]. Mice transplanted with control PP clusters, on the contrary, were not able to produce C-peptide in response to glucose 8 weeks post-transplantation (-AT7867 (t=0): 21.3 pM [IQR 19.7-22.4 pM]; -AT7867 (t=60): 19.3 pM [IQR 12.8-19.55 pM]; *p=0.8682*). However, at 12 weeks post-transplantation, mice transplanted with AT7867 treated PP clusters or control PP clusters had similar IPGTT profiles to naïve mice (**Figure 5L**). Measurement of AUC showed no significant difference between mice transplanted with AT7867 treated PP clusters or control PP clusters and naïve mice (-AT7867: 1884 pM [IQR1744-2024 pM]; *p=0.2352*; +AT7867: 1714 pM [IQR 1438-1990 pM]; *p=0.7043*; **Figure 5M**). 12 weeks post-transplantation, mice transplanted with AT7867 treated PP clusters or control PP clusters had similar C-peptide levels at time 0 (-AT7867: 120.6 pM [IQR 89.3-126.3 pM]; +AT7867: 117.2 [IQR 77.5-150.4 pM]); *p=0.9988*; **Figure 5N**) and showed similar glucose responsive C-peptide production 60 minutes after glucose administration (-AT7867: 213.2 pM [IQR 195.3-233.4 pM]; +AT7867: 248.3 pM [IQR 228.5-350.2 pM]; *p=0.1456*; **Figure 5M**). Histological assessment of the graft confirmed that AT7867 treatment did not hamper the *in vivo* differentiation into mature monohormonal insulin or glucagon secreting cells (**Figure 5O**).

## 3 Discussion

Our results show that addition of AT7867 during PP differentiation significantly increases the proportion of PDX1^+^NKX6.1^+^GP2^+^ cells without altering the total cell yield or proliferation of PPs. These results in combination with significant upregulation of genes associated with PP and PEP commitment and significant downregulation of pluripotency genes suggest that AT7867 induces differentiation of PP cells rather than proliferation. Furthermore, our results demonstrate that high purity of PP cells measured as >90% of PDX1^+^NKX6.1^+^GP2^+^ results in accelerated diabetes reversal following transplant and *in vivo* maturation.

The presence of uncommitted cells remains the major obstacle for clinical translation of SC-islets^36–39^. For this reason, strategies to enrich the pancreatic endocrine population and remove off-target cells, including methods involving chemical^40–44^, physical^3,5,39,45^ and/or genetic manipulation^46–48^, are becoming a focus for intense investigation. Key to pancreatic endocrine cells is the generation of pure PP cells. Previous studies have identified GP2 as a highly specific marker for PP cells capable to differentiate into insulin secreting cells *in vivo*^39,49^. As such, sorting of GP2^+^ cells followed by transplantation of 76% GP2^+^ PPs has previously been described as a method to eliminate contaminating off-target cells and the associated risk of teratoma formation post-transplantation^39^. However, sorting of GP2^+^ cells resulted in significant cell loss with a recovery of only 16% of the cells^39^, hence, making this strategy inappropriate for large-scale manufacturing for clinical translation. Optimization of the differentiation protocol to generate homogenous population is a more cost effective and scalable approach. Herein, we propose a scalable chemical-based approach using the small molecule AT7867 to generate PP populations with >90% PDX1^+^GP2^+^ that ultimately give rise to functional β-cells *in vivo*.

AT7867 is a potent AKT and p70 S6 kinase inhibitor used to slow the progression of tumor growth by inducing G2/M phase arrest and cell apoptosis in cancer stem cells^50,51^. AT7867 has also been described to induce the proliferation of PDX1^+^ cells; furthermore, Kimura et al described that the increased cell density as a result of AT7867 mediated proliferation triggered the upregulation of PDX1 and NKX6.1in PP cells^33^. However, our results suggest that the increased number of PDX1^+^NKX6.1^+^ is a result of improved differentiation rather than proliferation as suggested by upregulation of genes associated with acquisition of pancreatic progenitor state and commitment to pancreatic endocrine lineage and the downregulation of genes involved with the establishment and maintenance of pluripotency ^39,52^. Improved differentiation resulted in PPs that had accelerated maturation into insulin secreting cells *in vivo* and consequently, enhanced glucose-stimulated insulin response, and faster diabetes reversal. Altogether, our data shows that addition of small molecule AT7867 results in the generation of a homogeneous population of PP cells able to undergo accelerated endocrine differentiation *in vivo*.

Several studies have reported *in vitro* differentiation of PP cells into SC-islets followed by diabetes reversal upon transplantation in mice, but to date, no studies have reported the presence, (or absence) of teratomas upon transplantation of SC-islets^7,9,37,40,41,45^. Further *in vitro* differentiation of PP cells into endocrine cells could be a potentially safer alternative for clinical translation due to the decreased proliferation rate of cells upon endocrine commitment^2,53–55^. However, the generation of pure PP populations would remain a limiting factor to prevent teratoma formation. Furthermore, survival of β-cells upon transplantation might be challenging due to their high oxygen consumption rate, which would hamper their survival, resulting in increased number of endocrine cells required for transplantation compared to PP cells. In addition, the lower differentiation yield associated with endocrine differentiation is a critical limiting factor for scalability for clinical translation.

*In vitro* generation of SC-islets or PP cells provides a unique opportunity to deliver adequate islet or cell progenitor masses to ensure achievement of long-term insulin independence (i.e., of >20,000 IEQ/kg). Assuming a 50% cell loss throughout differentiation, we estimate that the generation and differentiation of up to 10^9^ iPSCs per patient would be required. Optimization of the differentiation protocol to minimize cell loss and increase yield is essential and generation of pure populations, rather than sorting of the desired population and potential cell loss of >80%^39^, is likely more cost effective for clinical translation. While planar culture conditions support scalability through the addition of more plates or flasks, suspension culture using VWB enables scalability into lager culture vessel formats^56^. For this reason, it is encouraging that our protocol was transferable to suspension culture using 0.1L VWB, showcasing potential to generate clinically relevant cell masses.

The outcomes of this study should be contextualized within specific limitations. Addition of AT7867 to this protocol has only been replicated with three healthy human donor-derived iPSC lines generated with Sendai virus mediated transfection of Yamanaka factors into PBMCs. Differentiation efficiency might vary based on the source of cell and the method used for reprogramming, as well as patient related factors including age, sex, or comorbidities. Replicating this protocol using iPSC lines from people with type 1 diabetes and/or other comorbidities will be essential to continue advancing the use of autologous iPSC-derived cellular replacement therapies. Importantly, our results remain limited to mouse models with renal subcapsular transplant; these results might not be translatable to other implant sites in rodents and/or human. Furthermore, long-term transplants to evaluate the safety of the transplanted cells would be required prior to clinical translation, and in-human safety and efficacy data would still be required to confirm these promising results. Furthermore, scalability of this protocol through differentiation using 0.5 L, 3 L or 15 L VWB remains to be tested. However, it is worth mentioning that the scalability of VWB to 0.5 L has been demonstrated for iPSC expansion and hence we do not expect any pitfalls in that aspect.

Despite these limitations, we present AT7867 as a novel small molecule that, when added during differentiation, improves *in vitro* differentiation efficiency of human iPSCs into PP cells. Following renal subcapsular transplant, AT7867 PPs are capable of *in vivo* maturation into monohormonal functional cells with accelerated diabetes reversal compared to controls. The potential to scale up this protocol using VWB represents a step towards the clinical translation of pluripotent stem cell-derived cellular therapeutics for the treatment of diabetes.

## 4 Experimental procedures

### 4.1 Resource availability

#### Corresponding author

Further information and requests for resources and reagents should be directed to and will be fulfilled by the corresponding author AM James Shapiro.

#### Materials availability

This study did not generate new unique reagents.

#### Data and code availability

All data reported in this paper will be shared by the lead contact upon request. This paper does not report original code. Any additional information required to reanalyse the data reported in this paper is available from the lead contact upon reasonable request.

### 4.2 Experimental model and subject details

Blood sample donors provided written consent for cell reprogramming, differentiation and result disclosure. This study and its methods have been approved by the Stem Cell Oversight Committee (SCOC), Canada and the University of Alberta Institutional Health Research Ethics Board (PRO00084032). Animal protocols were conducted in accordance with the Canadian Council on Animal Care Guidelines and Policies and have been approved by the Animal Care and Use Committee (Health Sciences) at the University of Alberta.

### 4.3 Cell culture

Three human iPSC lines generated from peripheral blood mononuclear cells (PBMCs) of healthy donors (patient demographics in **Supplementary Material** Error! Reference source not found.**S5**) were used. iPSC lines were generated through Sendai virus mediated PBMC transfection^57^. iPSCs were cultured on recombinant human VTN (rhVTN) coated 60mm plates in StemFlex media (Stem Cell Technologies, cat. A3349401) and passaged using CTS EDTA Versene Solution (Fisher Scientific, cat. A4239101) supplemented with 10 μM Rho-kinase inhibitor (RockI; Y-27632 STEMCELL Technologies, cat. 72304). iPSCs were seeded at a density of 6×10^4^ cells/cm^2^ and expanded for 3 days to achieve 80% confluency prior to differentiation. Confluency was monitored with the ECHO Rebel inverted microscope (ECHO). For expansion in VWB 3.6×10^4^ live cells/ mL were seeded into a 0.1 L Vertical Wheel® Bioreactor using 55 mL of StemFlex media supplemented with 10 μM RockI and expanded for 5 days prior to differentiation at 60 rotations per minute (rpm). Cells were counted and viability was assessed using the Thermo Fisher Scientific Invitrogen Countess II AMQAX1000 Cell Counter.

Differentiation into PPs was carried out using a published four stage protocol with modifications^35,58^. iPSCs were cultured for 4 days using STEMdiff™ Definitive Endoderm Differentiation Kit (Cat. No. 05110, STEMCELL Technologies). Media was replaced with RPMI 1640 medium, GlutaMAX supplemented (Cat. No. 61870-036, Thermo Fisher Scientific) supplemented with 1% (v/v) B-27 Serum-Free Supplement (50x) (Cat. No. 17504-001, Thermo Fisher Scientific) and 50 ng/ml KGF (Cat. No. 251-KG-MTO, R&D System) for 2 days. From day 6 to day 8, media was changed to StableCell DMEM-High Glucose (Cat. No. D0822-500ML, Sigma) supplemented with 1% (v/v) B-27, 0.25 μM KAAD-Cyclopamine (Cat. No. 239804, EMD Millipore), 2 μM Retinoic acid (Cat. No. 0695, Tocris Bioscience) and 0.25 μM LDN193189 (Cat. No. 04-0074, Stemgent). Cells were then cultured in DMEM supplemented with 1% (v/v) B-27, 50 ng/ml EGF (Cat. No. 236-EG, R&D System), 25 ng/ml FGF7 and 1 μM AT7867 (Cat. No. 7001, Tocris) for 4 days. Media changes were performed by aspirating the used media in plates or by allowing the clusters to gravity settle before the removal of the supernatant followed, in both cases, by addition of freshly prepared media. Morphology of the cells and clusters was assessed with the ECHO Rebel inverted microscope.

### 4.4 Flow cytometry

Cells differentiated in plates were lifted using CTS EDTA Versene solution, while clusters were dissociated with Accutase. Cells were strained through a 40 µm strainer prior to fixation in 4% paraformaldehyde (PFA). Cells were permeabilized using Cytofix/Cytoperm (Cat. No. 554714, BD Biosciences) for 20LJminutes on ice followed by 2 washes with 1x Perm/ Wash buffer (Cat. No. 554714, BD Biosciences). Primary antibodies were incubated for 1 hour (hr) on ice and secondary antibodies for 30 minutes on ice (dilutions in **Table S6**). Cells were resuspended in fluorescence-activated cell sorting (FACS) buffer (2% FCS, 2 mM EDTA in DPBS). For DNA content measurement, cells were stained with LIVE/DEAD Fixable Near-IR (Cat. No. L34975, Thermo Fisher Scientific) prior to fixation in 4% PFA. Fixed cells were then permeabilized as above and stained with 1 μM Hoechst 33342 (Cat. No. H1399, Thermo Fisher Scientific) for 30 minutes at room temperature in the dark. Cells were washed with 1x Perm/ Wash buffer and resuspended in FACS buffer prior to data acquisition. Data were acquired using the CytoFLEX S flow cytometer and analysed using the CytExpert software (Beckman Coulter).

### 4.5 Western Blotting

Cells were lysed in a buffer containing 50 mM Tris (pH 8.0), 150 mM NaCl, and 1% *v*/*v* Triton X-100 and supplemented with protease inhibitor tablets (Cat. No. A32955, Thermo Fisher Scientific). Protein concentrations were determined by BCA protein assay. 30 μg of heat-denatured proteins were run on Mini-PROTEAN TGX Precast Gels, 10% Novex (Cat. No. 456-1033, BioRad) and electrically transferred to nitrocellulose membranes. After blocking for 1 h at room temperature with 1% BSA, membranes were incubated overnight at 4 °C with primary antibodies. PDX1 (Cat. No. AF2419, R&D System) and KI67 (Cat. No. ab15580, Abcam) were diluted 1:1000 on blocking buffer. The next day, membranes were incubated with horseradish-peroxidase-linked secondary antibodies diluted 1:5000 followed by exposure to Clarity Western ECL Substrate (Cat. No. 170-5060, BioRad) and film development. Human β-ACTIN (Cat. No. AM4302, Invitrogen) was used as loading control.

### 4.6 Immunohistochemistry

iPSCs were grown and differentiated into geltrex (Cat. No. A1413301, Thermo Fisher Scientific) coated coverslips and fixed in 4% PFA for 20 minutes at room temperature. Tissue cross-sections were deparaffinized and rehydrated and subjected to antigen retrieval using citrate buffer (0.0126 M citric acid, Cat. No. C-0759, Sigma; 0.0874 M sodium citrate, Cat. No. S-4641, Sigma; pH 6.0) for a total of 20 minutes. Coverslips and tissue cross-sections were blocked and permeabilized with 5% normal donkey serum (Cat. No. S30-M, Sigma) in FoxP3 permeabilization buffer (Cat. No. 421402, Biolegend) for 1 hr at room temperature and incubated with primary antibodies overnight at 4 °C. Secondary antibodies were incubated for 30 minutes at room temperature in the dark followed by DAPI (Cat. No. D1306, Sigma) staining for 4 minutes at room temperature. Antibodies and concentrations used are listed in **Table S5**. Slides were visualized using the Zeiss Observer Z1 inverted fluorescence microscope and images were processed using Zeiss software and analyzed using QuPath ^59^.

### 4.7 qRT-PCR

Custom designed gene TaqMan Low Density Array Cards were used as per manufacturer instructions (Cat. No. 4342253, Thermo Fisher Scientific); gene array set ups are described in **Table S7** and **S8**. Data acquisition was performed in QuantStudio 12K Flex Real-Time PCR system. Samples were analysed using GAPDH as reference for data normalization. Data was analyzed and represented as a heatmap, 2^(-ΔΔCT)^ or Volcano plots using GraphBio^60^. GraphPad Prism version 9.3.1 or VolcanoSer^61^.

### 4.8 Diabetic induction and transplantation

Five days prior to transplantation, diabetes was induced by intraperitoneal (IP) injection of 75 mg/kg of streptozotocin (STZ; Cat. No. 572201, Millipore Sigma) in acetate buffer, pH 4.5. STZ IP injections were repeated for up to 4 days until SCID beige mice 10 to 14-week-old and balanced for sex were considered diabetic following a non-fasting blood glucose measurement of ≥15.0 mmol/L on two consecutive days. Only animals meeting this inclusion criterion were selected for transplantation. 1500 PP clusters were transplanted under the kidney capsule^62^ and an erodible insulin pellet (Cat. no. As-1-L, LinShin Canada, LinBit, 0.1U/24hr/implant) was implanted subcutaneously to maintain animal health over a 30-day period. Mice were anesthetized with 5% isoflurane. Buprenorphine (0.1 mg/kg subcutaneous) was administered for post-operative analgesia. Mice were assessed daily for humane endpoints.

On post-operative day 90, non-recovery nephrectomy was performed; mice were euthanized under anesthesia by clipping the heart. Kidney cross-sections were fixed in 10% formalin, and paraffinized. 8 µm sections were prepared for immunohistochemistry as above.

### 4.9 Assessment of glycaemic control

Glycemic control was assessed using non-fasting blood glucose measurements (mM) three times a week after transplantation using a portable glucometer (OneTouch Ultra 2, LifeScan).

Intraperitoneal glucose tolerance tests (IPGTT) were conducted at 8- and 12-weeks post-transplant. Animals were fasted overnight before receiving 3 mg/g of weight via IP. Blood glucose levels were monitored prior to IP injection and 15, 30, 60, 90 and 120 minutes post-injection. blood samples were collected prior to IP injection and 60 minutes post-injection to measure human C-peptide content with enzyme-linked immunosorbent assay (Cat. No. 10-1136-01, Mercodia).

### 4.10 Statistical Analysis

Normality testing was performed using the D’Agostino-Pearson normality test, which determined the need for non-parametric testing. Between group comparisons were carried out using the non-parametric Mann–Whitney U test or Kruskal–Wallis test. Two-way Anova was used to compare time courses. The alpha value was set at 0.05 but was modified *post hoc* to 0.01 for volcano plot evaluation of transcriptomic data to better display key gene expression changes. Continuous values are presented as medians with interquartile ranges (IQR), and with discrete values presented as absolute values with percentages. All statistical analysis was completed using GraphPad Prism version 9.3.1 for Mac, GraphPad Software, www.graphpad.com.

## Acknowledgements

AMJS is supported through a Canada Research Chair (Tier 1) in Regenerative Medicine and Transplant Surgery, and through grant support from the Juvenile Diabetes Research Foundation, Diabetes Canada, the Canadian Donation and Transplant Research Program, the Diabetes Research Institute Foundation of Canada, the Alberta Diabetes Foundation, and the Canadian Stem Cell Network. Braulio A. Marfil–Garza is supported by the CHRISTUS Excellence and Innovation Center.

## Author Contributions

NCG participated in study conceptualization, data curation, formal analysis, investigation, methodology, writing of the original draft, and final draft review and editing. KV and ND participated in study conceptualization, data curation, investigation, methodology, writing of the original draft, and final draft review and editing. RP, MBG and HR participated in data curation, investigation, methodology, and final draft review and editing. AMJS participated in study conceptualization, formal analysis, methodology, funding acquisition, project administration, supervision, and final draft review and editing. AMJS supervised this project’s work, is responsible for the data within the study, has ensured that all authorship is granted appropriately with all disclosures identified and has ensured all authors have approved the work.

## References

1. Shapiro, A. M. J. et al. Islet transplantation in seven patients with type 1 diabetes mellitus using a glucocorticoid-free immunosuppressive regimen. N Engl J Med 343, 230–238 (2000).

2. Balboa, D. et al. Functional, metabolic and transcriptional maturation of human pancreatic islets derived from stem cells. Nat Biotechnol 40, 1042–1055 (2022).

3. Parent, A. V et al. Stem Cell Reports Resource Development of a scalable method to isolate subsets of stem cell-derived pancreatic islet cells. Stem Cell Reports 17, 979–992 (2022).

4. Verhoeff, K. et al. Optimizing Generation of Stem Cell-Derived Islet Cells. Stem Cell Reviews and Reports 2022 18:8 18, 2683–2698 (2022).

5. Nair, G. G. et al. Recapitulating endocrine cell clustering in culture promotes maturation of human stem-cell-derived β cells. Nature Cell Biology 2019 21:2 21, 263–274 (2019).

6. Kroon, E. et al. Pancreatic endoderm derived from human embryonic stem cells generates glucose-responsive insulin-secreting cells in vivo. Nature Biotechnology 2008 26:4 26, 443–452 (2008).

7. Rezania, A. et al. Reversal of diabetes with insulin-producing cells derived in vitro from human pluripotent stem cells. Nature Biotechnology 2014 32:11 32, 1121–1133 (2014).

8. Velazco-Cruz, L. et al. Stem Cell Reports Article Acquisition of Dynamic Function in Human Stem Cell-Derived b Cells. Stem Cell Reports 12, 351–365 (2019).

9. Pagliuca, F. W. et al. Generation of functional human pancreatic β cells in vitro. Cell 159, 428–439 (2014).

10. Millman, J. R. et al. Generation of stem cell-derived β-cells from patients with type 1 diabetes. Nature Communications 2016 7:1 7, 1–9 (2016).

11. Hogrebe, N. J., Augsornworawat, P., Maxwell, K. G., Velazco-Cruz, L. & Millman, J. R. Targeting the cytoskeleton to direct pancreatic differentiation of human pluripotent stem cells. Nat Biotechnol 38, 460–470 (2020).

12. Hogrebe, N. J., Maxwell, K. G., Augsornworawat, P. & Millman, J. R. Generation of insulin-producing pancreatic β cells from multiple human stem cell lines. Nat Protoc 16, 4109– 4143 (2021).

13. Aghazadeh, Y. et al. GP2-enriched pancreatic progenitors give rise to functional beta cells in vivo and eliminate the risk of teratoma formation. Stem Cell Reports 17, 964–978 (2022).

14. Rostovskaya, M., Bredenkamp, N. & Smith, A. Towards consistent generation of pancreatic lineage progenitors from human pluripotent stem cells. Philosophical Transactions of the Royal Society B: Biological Sciences 370, (2015).

15. Bastidas-Ponce, A. et al. Comprehensive single cell mRNA profiling reveals a detailed roadmap for pancreatic endocrinogenesis. Development (Cambridge) 146, (2019).

16. Veres, A. et al. Charting cellular identity during human in vitro β-cell differentiation. Nature 2019 569:7756 569, 368–373 (2019).

17. Wesolowska-Andersen, A. et al. Deep learning models predict regulatory variants in pancreatic islets and refine type 2 diabetes association signals. Elife 9, (2020).

18. Sander, N. et al. Homeobox gene Nkx6.1 lies downstream of Nkx2.2 in the major pathway of beta-cell formation in the pancreas. Development 127, 5533–5540 (2000).

19. Nelson, S. B., Schaffer, A. E. & Sander, M. The transcription factors Nkx6.1 and Nkx6.2 possess equivalent activities in promoting beta-cell fate specification in Pdx1+ pancreatic progenitor cells. Development 134, 2491–2500 (2007).

20. Rukstalis, J. M. & Habener, J. F. Neurogenin3: a master regulator of pancreatic islet differentiation and regeneration. Islets 1, 177–184 (2009).

21. Ramond, C. et al. Reconstructing human pancreatic differentiation by mapping specific cell populations during development. Elife 6, (2017).

22. Krentz, N. A. J. et al. Single-Cell Transcriptome Profiling of Mouse and hESC-Derived Pancreatic Progenitors. Stem Cell Reports 11, 1551–1564 (2018).

23. Wang, H., Brun, T., Kataoka, K., Sharma, A. J. & Wollheim, C. B. MafA controls Genes implicated in Insulin Biosynthesis and Secretion. Diabetologia 50, 348 (2007).

24. Augsornworawat, P., Maxwell, K. G., Velazco-Cruz, L., Millman Correspondence, J. R. & Millman, J. R. Single-Cell Transcriptome Profiling Reveals b Cell Maturation in Stem Cell-Derived Islets after Transplantation. doi:10.1016/j.celrep.2020.108067.

25. Sui, L., et al. Reduced replication fork speed promotes pancreatic endocrine differentiation and controls graft size. JCI Insight 6, (2021).

26. Ben-David, U., Nudel, N. & Benvenisty, N. Immunologic and chemical targeting of the tight-junction protein Claudin-6 eliminates tumorigenic human pluripotent stem cells. Nat Commun 4, 1992 (2013).

27. Verhoeff, K. et al. Optimizing Generation of Stem Cell-Derived Islet Cells. Stem Cell Reviews and Reports 2022 18:8 18, 2683–2698 (2022).

28. Zhang, Y. et al. Nicotinamide promotes pancreatic differentiation through the dual inhibition of CK1 and ROCK kinases in human embryonic stem cells. Stem Cell Res Ther 12, 1–12 (2021).

29. Nostro, M. C. et al. Efficient Generation of NKX6-1+ Pancreatic Progenitors from Multiple Human Pluripotent Stem Cell Lines. Stem Cell Reports 4, 591 (2015).

30. Rezania, A. et al. Maturation of human embryonic stem cell-derived pancreatic progenitors into functional islets capable of treating pre-existing diabetes in mice. Diabetes 61, 2016–2029 (2012).

31. Liu, H. et al. Chemical combinations potentiate human pluripotent stem cell-derived 3D pancreatic progenitor clusters toward functional β cells. Nature Communications 2021 12:1 12, 1–10 (2021).

32. Jiang, Y. et al. Stem Cell Reports Resource Generation of pancreatic progenitors from human pluripotent stem cells by small molecules. (2021) doi:10.1016/j.stemcr.2021.07.021.

33. Kimura, A., et al. Small molecule AT7867 proliferates PDX1-expressing pancreatic progenitor cells derived from human pluripotent stem cells. Stem Cell Res 24, 61–68 (2017).

34. Elghazi, L. et al. Regulation of Pancreas Plasticity and Malignant Transformation by Akt Signaling. Gastroenterology 136, 1091 (2009).

35. Sui, L., Leibel, R. L. & Egli, D. Pancreatic Beta Cell Differentiation From Human Pluripotent Stem Cells. Curr Protoc Hum Genet 99, (2018).

36. Kelly, O. G. et al. Cell-surface markers for the isolation of pancreatic cell types derived from human embryonic stem cells. Nat Biotechnol 29, 750–756 (2011).

37. Rezania, A. et al. Maturation of human embryonic stem cell-derived pancreatic progenitors into functional islets capable of treating pre-existing diabetes in mice. Diabetes 61, 2016–2029 (2012).

38. Kroon, E. et al. Pancreatic endoderm derived from human embryonic stem cells generates glucose-responsive insulin-secreting cells in vivo. Nature Biotechnology 2008 26:4 26, 443–452 (2008).

39. Aghazadeh, Y. et al. GP2-enriched pancreatic progenitors give rise to functional beta cells in vivo and eliminate the risk of teratoma formation. Stem Cell Rep. 17, 964–978 (2022).

40. Hogrebe, N. J., Augsornworawat, P., Maxwell, K. G., Velazco-Cruz, L. & Millman, J. R. Targeting the cytoskeleton to direct pancreatic differentiation of human pluripotent stem cells. Nature Biotechnology 2020 38:4 38, 460–470 (2020).

41. Velazco-Cruz, L. et al. Acquisition of Dynamic Function in Human Stem Cell-Derived β Cells. Stem Cell Reports 12, 351–365 (2019).

42. Lee, M. O. et al. Inhibition of pluripotent stem cell-derived teratoma formation by small molecules. Proc Natl Acad Sci U S A 110, E3281–E3290 (2013).

43. Ben-David, U. & Benvenisty, N. Chemical ablation of tumor-initiating human pluripotent stem cells. Nature Protocols 2014 9:3 9, 729–740 (2014).

44. Sui, L., et al. Reduced replication fork speed promotes pancreatic endocrine differentiation and controls graft size. JCI Insight 6, (2021).

45. Veres, A. et al. Charting cellular identity during human in vitro β-cell differentiation. Nature 2019 569:7756 569, 368–373 (2019).

46. Zhou, X. et al. Serial Activation of the Inducible Caspase 9 Safety Switch After Human Stem Cell Transplantation. Molecular Therapy 24, 823 (2016).

47. Schuldiner, M., Itskovitz-Eldor, J. & Benvenisty, N. Selective ablation of human embryonic stem cells expressing a ‘suicide’ gene. Stem Cells 21, 257–265 (2003).

48. Chen, F. et al. Suicide gene-mediated ablation of tumor-initiating mouse pluripotent stem cells. Biomaterials 34, 1701–1711 (2013).

49. Ameri, J. et al. Efficient Generation of Glucose-Responsive Beta Cells from Isolated GP2+ Human Pancreatic Progenitors. Cell Rep 19, 36–49 (2017).

50. Zhang, S., et al. AT7867 Inhibits Human Colorectal Cancer Cells via AKT-Dependent and AKT-Independent Mechanisms. PLoS One 12, (2017).

51. Li, Y., et al. AT7867 Inhibits the Growth of Colorectal Cancer Stem-Like Cells and Stemness by Regulating the Stem Cell Maintenance Factor Ascl2 and Akt Signaling. Stem Cells Int 2023, (2023).

52. Heller, S. et al. Transcriptional changes and the role of ONECUT1 in hPSC pancreatic differentiation. Commun Biol 4, (2021).

53. McDonald, E. et al. SOX9 regulates endocrine cell differentiation during human fetal pancreas development. Int J Biochem Cell Biol 44, 72–83 (2012).

54. Rosado-Olivieri, E. A., Aigha, I. I., Kenty, J. H. & Melton, D. A. Identification of a LIF-Responsive, Replication-Competent Subpopulation of Human β Cells. Cell Metab 31, 327–338.e6 (2020).

55. Oakie, A. & Nostro, M. C. Harnessing Proliferation for the Expansion of Stem Cell-Derived Pancreatic Cells: Advantages and Limitations. Front Endocrinol (Lausanne) 12, 91 (2021).

56. Dang, T. et al. Computational fluid dynamic characterization of vertical-wheel bioreactors used for effective scale-up of human induced pluripotent stem cell aggregate culture. Can J Chem Eng 99, 2536–2553 (2021).

57. Rim, Y., Nam, Y. & Ju, J. Induced Pluripotent Stem Cell Generation from Blood Cells Using Sendai Virus and Centrifugation. J Vis Exp 2016, 54650 (2016).

58. Sui, L., et al. Reduced replication fork speed promotes pancreatic endocrine differentiation and controls graft size. JCI Insight 6, (2021).

59. Bankhead, P. et al. QuPath: Open source software for digital pathology image analysis. Scientific Reports 2017 7:1 7, 1–7 (2017).

60. Zhao, T. & Wang, Z. GraphBio: A shiny web app to easily perform popular visualization analysis for omics data. Front Genet 13, 2265 (2022).

61. Goedhart, J. & Luijsterburg, M. S. VolcaNoseR is a web app for creating, exploring, labeling and sharing volcano plots. Scientific Reports 2020 10:1 10, 1–5 (2020).

62. Szot, G. L., Koudria, P. & Bluestone, J. A. Transplantation of Pancreatic Islets Into the Kidney Capsule of Diabetic Mice. J Vis Exp 9, (2007).

