## Supplementary material for "AT7867 promotes pancreatic progenitor differentiation of human iPSCs and accelerates diabetes reversal"

**Supplementary Figures**

**Figure S1. Differentiation of 3 independent iPSC lines using AT7867.** A) Quantification of flow cytometry analysis at iPSC stage, B) stage 1, C) stage 2, D) stage 3 and E) stage 4 of three independent iPSC lines from 6 independent experiments.

**Figure S2. AT7867 does not induce the proliferation of pancreatic progenitor cells. A)** Representative immunohistochemistry of PDX1 and KI67 expression. B) Representative flow cytometry gating of KI67^+^ cells from day 8 to day 12 of differentiation.

**Figure S3. Fold change of genes associated to pancreatic endocrine differentiation.**

**Figure S4. Fold change of genes associated to pluripotency and non-endocrine pancreatic differentiation.**

**Figure S5. iPSC differentiation into Vertical Wheel® bioreactors.** A) Quantification of flow cytometry analysis at iPSC stage, B) stage 1, C) stage 2 and D) stage 3 of iPSCs from 6 independent experiments cultured and differentiated in vertical Wheel® bioreactors. E) Cluster size and quantification of flow cytometry analysis at stage 4 with and without AT7867 of iPSCs from 6 independent experiments cultured in bioreactors.

**Supplementary Tables**

**Table S1. Percentage of proliferative cells (KI67^+^) for control and AT7867 treated cells from day 8 to day 12.** Between group comparisons were carried out using the non-parametric Mann-Whitney U test.

|  | **- AT7867** | **+ AT7867** | **p value** |
| --- | --- | --- | --- |
| **Day 8** | 82.43% (IQR 81.69% - 85.44%) | | - |
| **Day 9** | 82.73% (IQR 81.16% - 85.46%) | 88.52% (IQR 86.20% - 89.69%) | *0.7253* |
| **Day 10** | 87.55% (IQR 87.29% - 88.64%) | 89.25% (IQR 85.65% - 91.12%) | *>0.9999* |
| **Day 11** | 75.66% (IQR 72.02% - 83.51%) | 82.00% (IQR 79.70% - 84.39%) | *0.7388* |
| **Day 12** | 78.42% (IQR 75.52% - 89.29%) | 81.27% (IQR 81.20% - 83.57%) | *>0.9999* |

**Table S2. Percentage of cells in G0/G1, S and G2/M phases for control and AT7867 treated cells from day 8 to day 12.** Between group comparisons were carried out using the non-parametric Mann-Whitney U test.

|  |  | **Day 8** | **Day 9** | **Day 10** | **Day 11** | **Day 12** |
| --- | --- | --- | --- | --- | --- | --- |
| **G0/G1 phase** | **-AT7867** | 22.85% (IQR 12.90% - 37.14%) | 14.34% (IQR 11.34% - 22.88%) | 24.42% (IQR 19.24% - 35.19%) | 16.53% (IQR 14.09% - 24.34%) | 28.05% (IQR 21.52% - 38.93%) |
|  | **+AT7867** |  | 21.40% (IQR 10.01% - 37.19%) | 31.93% (IQR 19.29% - 36.35%) | 16.38% (IQR 15.09% - 27.95%) | 2021% (IQR 17.35% - 21.26%) |
|  | **p value** | - | *0.9866* | *>0.9999* | *>0.9999* | *0.8844* |
| **S phase** | **-AT7867** | 47.82% (IQR 43.95% - 51.81%) | 44.97% (IQR 43.45% - 49.66%) | 47.29% (IQR 44.16% - 47.40%) | 48.19% (IQR 47.98% - 49.46%) | 46.94% (IQR 38.21% - 50.06%) |
|  | **+AT7867** |  | 39.04% (IQR 38.13% - 42.43%) | 47.36% (IQR 44.60% - 52.58%) | 49.33% (IQR 46.18% - 50.44%) | 46.90% (IQR 42.26% - 48.29%) |
|  | **p value** | - | *0.4346* | *0.9984* | *>0.9999* | *>0.9999* |
| **G2/M** | **-AT7867** | 30.59% (IQR 17.88% - 38.61%) | 42.21% (27.46% - 43.69%) | 20.14% (IQR 18.74% - 27.53%) | 33.44% (IQR 27.30% - 37.01%) | 46.94% (IQR 38.21% - 50.06%) |
|  | **+AT7867** |  | 40.47% (IQR 23.77% - 47.56%) | 27.38% (IQR 17.45% - 35.91%) | 32.59% (IQR 25.22% - 34.76%) | 46.90% (42.26% - 48.29%) |
|  | **p value** | - | *>0.9999* | *0.997* | *>0.9999* | *>0.9999* |

**Table S3. Median, interquartile range and p value for genes represented in Figure 3B-K and Figure 4B-K.**

| **Gene** | **- AT7867** | **+ AT7867** | **p value** |
| --- | --- | --- | --- |
| ***ALPL*** | 0.61 (IQR 0.47 – 0.78) | 0.12 (IQR 0.08 – 0.13) | *0.0029* |
| ***ARX*** | 26.48 (IQR 19.31 – 31.11) | 281.4 (IQR 173.9 – 283.2) | *0.0037* |
| ***FGF4*** | 1.78 (IQR 1.06 – 2.10) | 0.30 (IQR 0.19 – 0.36) | *0.0117* |
| ***GDF3*** | 0.01 (IQR 0.01 – 0.02) | 0.10 (IQR 0.10 – 0.20) | *0.0262* |
| ***GP2*** | 829.4 (IQR 766.4 – 964.6) | 1782 (IQR 1584 – 2085) | *0.0007* |
| ***HNF4A*** | 9183 (IQR 6406 – 10791) | 58756 (IQR47226 – 62188) | *< 0.0001* |
| ***KIT*** | 2.04 (IQR 1.80 – 2.11) | 0.56 (IQR 0.47 – 0.57) | *< 0.0001* |
| ***KRT19*** | 2.52 (IQR 2.17 – 3.89) | 8.01 (IQR 7.64 – 10.97) | *0.0019* |
| ***LIN28A*** | 0.31 (IQR 0.28 – 0.37) | 0.14 (IQR 0.09 – 0.18) | *0.0356* |
| ***MYC*** | 0.09 (IQR 0.08 – 0.13) | 0.05 (IQR 0.03 – 0.06) | *0.0482* |
| ***NEUROD1*** | 10.39 (IQR 9.71 – 11.76) | 81.56 (IQR 81.39 – 83.33) | *< 0.0001* |
| ***NEUROG3*** | 1.35 (IQR 1.34 – 1.96) | 2.77 (IQR 2.60 – 3.44) | *0.0054* |
| ***NKX2.2*** | 111.4 (IQR 98.58 – 144.7) | 1148 (IQR 915.5 – 1511) | *0.0007* |
| ***NKX6.1*** | 153.6 (IQR135.1 – 194.3) | 9599 (IQR 7652 – 10238) | *< 0.0001* |
| ***ONECUT1*** | 5.90 (IQR 5.34 – 12.52) | 216.5 (IQR 159.1 – 219.7) | *< 0.0001* |
| ***PAX4*** | 8.29 (IQR 4.09 – 12.35) | 199.8 (IQR 125.2 – 199.8) | *0.0014* |
| ***PDX1*** | 621.7 (IQR 490.8 – 762.1) | 7233 (IQR 4752 – 7296) | *0.0004* |
| ***PODXL*** | 0.08 (IQR 0.07 – 0.12) | 0.01 (IQR 0.01 – 0.02) | *0.0057* |
| ***SOX9*** | 5.90 (IQR 3.20 – 7.32) | 21.78 (IQR 17.35 – 24.59) | *0.0005* |
| ***TBX2*** | 280.1 (IQR 235.4 – 286.7) | 10.72 (IQR 7.93 – 12.68) | *< 0.0001* |

**Table S4. Fold change and significance of the transcripts of AT7867 treated PP cells vs. control PP cells.**

| **Gene** | **Fold change** | **Fold change (Log2)** | **p value** | **Significance** | **Manhattan distance** |
| --- | --- | --- | --- | --- | --- |
| ***TBX2*** | 0.03829 | -4.70690 | 0.00001 | 4.86939 | 9.57629 |
| ***SOX17*** | 0.09841 | -3.34510 | 0.00007 | 4.16042 | 7.50553 |
| ***PDX1*** | 11.63320 | 3.54018 | 0.00013 | 3.89273 | 7.43291 |
| ***ONECUT1*** | 36.64686 | 5.19562 | 0.00017 | 3.77148 | 8.96710 |
| ***SOX9*** | 17.99033 | 4.16915 | 0.00019 | 3.71606 | 7.88521 |
| ***HNF4A*** | 6.39811 | 2.67764 | 0.00022 | 3.65661 | 6.33426 |
| ***CHGA*** | 8.10018 | 3.01795 | 0.00023 | 3.64314 | 6.66109 |
| ***ARX*** | 10.62839 | 3.40985 | 0.00023 | 3.64178 | 7.05164 |
| ***PODXL*** | 0.21594 | -2.21132 | 0.00028 | 3.54923 | 5.76055 |
| ***ISL1*** | 3.29592 | 1.72068 | 0.00048 | 3.32118 | 5.04186 |
| ***PAX4*** | 24.08147 | 4.58985 | 0.00053 | 3.27527 | 7.86513 |
| ***NKX6.1*** | 62.48049 | 5.96533 | 0.00058 | 3.23521 | 9.20054 |
| ***ALPL*** | 0.19705 | -2.34335 | 0.00058 | 3.23506 | 5.57841 |
| ***KIT*** | 0.27723 | -1.85085 | 0.00062 | 3.20747 | 5.05832 |
| ***FOXA2*** | 1.80166 | 0.84933 | 0.00083 | 3.07936 | 3.92868 |
| ***SYP*** | 0.13397 | -2.90004 | 0.00105 | 2.97755 | 5.87759 |
| ***NEUROD1*** | 7.83163 | 2.96931 | 0.00113 | 2.94843 | 5.91774 |
| ***CD274*** | 6.50845 | 2.70231 | 0.00122 | 2.91532 | 5.61763 |
| ***GDF3*** | 5.34383 | 2.41788 | 0.00145 | 2.83963 | 5.25750 |
| ***ITGA1*** | 1.86258 | 0.89730 | 0.00155 | 2.81051 | 3.70781 |
| ***NKX2.2*** | 10.30029 | 3.36461 | 0.00194 | 2.71139 | 6.07600 |
| ***GP2*** | 2.14889 | 1.10359 | 0.00228 | 2.64263 | 3.74622 |
| ***FGF10*** | 0.40256 | -1.31272 | 0.00258 | 2.58759 | 3.90030 |
| ***KRT19*** | 3.17850 | 1.66834 | 0.00263 | 2.58036 | 4.24870 |
| ***GCG*** | 16.14662 | 4.01316 | 0.00269 | 2.56951 | 6.58268 |
| ***HLA-B*** | 0.37265 | -1.42409 | 0.00321 | 2.49372 | 3.91782 |
| ***INS*** | 8.38826 | 3.06837 | 0.00353 | 2.45181 | 5.52018 |
| ***TERT*** | 0.47516 | -1.07351 | 0.00386 | 2.41296 | 3.48648 |
| ***HLA-A*** | 0.46016 | -1.11980 | 0.00493 | 2.30701 | 3.42681 |
| ***SLC2A4*** | 0.43697 | -1.19441 | 0.00518 | 2.28604 | 3.48045 |
| ***MAFB*** | 2.25075 | 1.17040 | 0.00560 | 2.25199 | 3.42239 |
| ***FGF4*** | 0.28564 | -1.80775 | 0.00666 | 2.17678 | 3.98454 |
| ***LIN28A*** | 0.44240 | -1.17658 | 0.00672 | 2.17250 | 3.34908 |
| ***PCSK1*** | 2.84132 | 1.50656 | 0.00695 | 2.15806 | 3.66462 |
| ***PAX6*** | 2.24026 | 1.16367 | 0.00888 | 2.05139 | 3.21506 |
| ***SST*** | 1.82262 | 0.86601 | 0.00911 | 2.04047 | 2.90648 |
| ***SLC30A8*** | 0.64451 | -0.63373 | 0.00987 | 2.00566 | 2.63939 |
| ***CPE*** | 0.74049 | -0.43344 | 0.01034 | 1.98540 | 2.41885 |
| ***SLC18A1*** | 4.11369 | 2.04043 | 0.01270 | 1.89634 | 3.93678 |
| ***IL6*** | 0.14601 | -2.77588 | 0.01308 | 1.88348 | 4.65936 |
| ***TSPAN1*** | 0.38164 | -1.38973 | 0.01337 | 1.87392 | 3.26365 |
| ***PODXL2*** | 0.53262 | -0.90882 | 0.01513 | 1.82006 | 2.72888 |
| ***SLC16A1*** | 0.56913 | -0.81317 | 0.01587 | 1.79930 | 2.61247 |
| ***MYC*** | 0.57315 | -0.80302 | 0.01650 | 1.78255 | 2.58556 |
| ***UCN3*** | 2.18716 | 1.12906 | 0.01950 | 1.70993 | 2.83898 |
| ***ABCC8*** | 2.62859 | 1.39429 | 0.02079 | 1.68218 | 3.07647 |
| ***FGF2*** | 0.62178 | -0.68554 | 0.03004 | 1.52231 | 2.20785 |
| ***FGFR1*** | 0.62351 | -0.68152 | 0.03040 | 1.51710 | 2.19861 |
| ***ABCG2*** | 0.54059 | -0.88738 | 0.03273 | 1.48507 | 2.37245 |
| ***PPY*** | 0.23494 | -2.08962 | 0.03499 | 1.45604 | 3.54566 |
| ***IL10*** | 0.24091 | -2.05341 | 0.04215 | 1.37522 | 3.42862 |
| ***G6PC2*** | 1.31275 | 0.39259 | 0.04440 | 1.35258 | 1.74517 |
| ***FOXO1*** | 0.67166 | -0.57420 | 0.04891 | 1.31061 | 1.88481 |
| ***ACVR2B*** | 0.53973 | -0.88970 | 0.05059 | 1.29592 | 2.18561 |
| ***HPRT1*** | 0.49750 | -1.00723 | 0.05186 | 1.28520 | 2.29243 |
| ***PCSK2*** | 0.60529 | -0.72431 | 0.06084 | 1.21581 | 1.94012 |
| ***POU5F1*** | 1.75093 | 0.80812 | 0.06949 | 1.15809 | 1.96621 |
| ***GATA4*** | 1.24752 | 0.31906 | 0.07737 | 1.11141 | 1.43047 |
| ***KCNK3*** | 1.80953 | 0.85561 | 0.07863 | 1.10441 | 1.96003 |
| ***NEUROG3*** | 2.04731 | 1.03373 | 0.08941 | 1.04861 | 2.08234 |
| ***IRX2*** | 1.80649 | 0.85319 | 0.10013 | 0.99942 | 1.85260 |
| ***B2M*** | 0.68789 | -0.53976 | 0.10473 | 0.97991 | 1.51967 |
| ***KCNK1*** | 1.22931 | 0.29785 | 0.12539 | 0.90174 | 1.19959 |
| ***MAFA*** | 1.25799 | 0.33113 | 0.12723 | 0.89542 | 1.22655 |
| ***CXCR4*** | 1.14274 | 0.19250 | 0.13491 | 0.86996 | 1.06245 |
| ***GCK*** | 2.60919 | 1.38360 | 0.13884 | 0.85750 | 2.24110 |
| ***KLF4*** | 1.69287 | 0.75947 | 0.15813 | 0.80100 | 1.56047 |
| ***TPBG*** | 4.69023 | 2.22966 | 0.23246 | 0.63365 | 2.86331 |
| ***SOX2*** | 1.05047 | 0.07103 | 0.25565 | 0.59235 | 0.66338 |
| ***IAPP*** | 1.57445 | 0.65485 | 0.27338 | 0.56324 | 1.21808 |
| ***CTLA4*** | 1.58891 | 0.66804 | 0.27557 | 0.55977 | 1.22781 |
| ***CDH1*** | 1.06884 | 0.09604 | 0.29235 | 0.53410 | 0.63014 |
| ***GLP1R*** | 0.76991 | -0.37724 | 0.30185 | 0.52020 | 0.89745 |
| ***CHGB*** | 0.98594 | -0.02042 | 0.32190 | 0.49228 | 0.51270 |
| ***CIITA*** | 0.96876 | -0.04579 | 0.33572 | 0.47403 | 0.51982 |
| ***ACVR1B*** | 1.17003 | 0.22654 | 0.37799 | 0.42252 | 0.64906 |
| ***HNF1A*** | 0.88242 | -0.18046 | 0.40150 | 0.39632 | 0.57678 |
| ***UTF1*** | 1.22226 | 0.28955 | 0.46878 | 0.32904 | 0.61858 |
| ***ZFP42*** | 1.22226 | 0.28955 | 0.46878 | 0.32904 | 0.61858 |
| ***ITGA6*** | 0.97940 | -0.03003 | 0.48731 | 0.31220 | 0.34223 |
| ***FUT4*** | 0.97771 | -0.03253 | 0.49485 | 0.30552 | 0.33805 |
| ***NANOG*** | 1.21305 | 0.27864 | 0.49834 | 0.30248 | 0.58111 |

**Table S5. Patient demographics used in this study.**

| **iPSC line** | **Age** | **Sex** | **Gender** | **Health status** |
| --- | --- | --- | --- | --- |
| **#1** | 53 | Female | Female | Healthy |
| **#2** | 43 | Female | Female | Healthy |
| **#4** | 29 | Male | Male | Healthy |

**Table S5. Antibodies and concentrations used for immunohistochemistry (IHC) and flow cytometry (FC).**
*All secondaries for immunohistochemistry were applied at a 1:250 concentration and all secondaries for flow cytometry were applied at a 1:500 concentration.

| **Epitope** | **Origin animal** | **Conjugate** | **Dilution** | **Supplier** | **Assay** |
| --- | --- | --- | --- | --- | --- |
| OCT4 | Mouse | BV421 | 1:100 | BD (565644) | FC |
| SSEA4 | Mouse | N/A | 1:100 | Invitrogen (MA1-021) | FC |
| SOX2 | Mouse | Fitc | 1:100 | Invitrogen (53-9811-82) | FC |
| NANOG | Mouse | Pe | 1:100 | Invitrogen (PA5-46891) | FC |
| CD184 | Mouse | BV421 | 1:100 | BD (562448) | FC |
| CD117 | Mouse | Fitc | 1:50 | Invitrogen (11-1178-42) | FC |
| SOX17 | Mouse | APC | 1:20 | R&D Systems (IC1924A) | FC |
| FOXA2 | Rabbit | N/A | 1:100 | Abcam (108422) | FC |
| PDX1 | Mouse | Fitc | 1:20 | BD (562274) | FC |
| PDX1 | Goat | N/A | 1:20 | R&D Systems (AF2419) | IHC |
| NKX6.1 | Mouse | N/A | 1:10 | DSHB (F55A10-c) | FC |
| CHGA | Mouse | AF405 | 1:50 | Novus (NBP2-33198AF405) | FC |
| GP2 | Mouse | AF405 | 1:50 | Novus (NBP3-08243AF405) | FC |
| KI67 | Mouse | PerCp-Cy5 | 1:50 | BD (561284) | FC |
| KI67 | Rabbit | N/A | 1:50 | Abcam (ab15580) | IHC |
| INSULIN | Guinea Pig | N/A | 1:500 | DAKO (A0564) | IHC |
| GLUCAGON | Mouse | N/A | 1:800 | Sigma (G2654) | IHC |
| Anti-goat | Donkey | Fitc | - | Thermo Fisher Scientific (A16000) | IHC |
| Anti-mouse | Donkey | AF647 | - | Invitrogen (A31571) | FC/ IHC |
| Anti-rabbit | Goat | AF594 | - | Invitrogen (cat. A11012) | IHC |
| Anti-guinea pig | Goat | Fitc | - | Invitrogen (cat. A11073) | FIHC |

**Table S6. Thermo Fisher TaqMan Micro Array configuration for the analysis of genes associated with islet differentiation.**

| **Assay ID** | **Gene** | **Gene Name(s)** | **Amplicon Length** |
| --- | --- | --- | --- |
| Hs01093752_m1 | *ABCC8* | ATP binding cassette subfamily C member 8 | 58 |
| Hs00292465_m1 | *ARX* | Aristaless related homeobox | 96 |
| Hs00900370_m1 | *CHGA* | Chromogranin A | 67 |
| Hs01084631_m1 | *CHGB* | Chromogranin B | 112 |
| Hs00175676_m1 | *CPE* | Carboxypeptidase E | 106 |
| Hs00607978_s1 | *CXCR4* | C-X-C motif chemokine receptor 4 | 153 |
| Hs00204257_m1 | *CD274* | CD274 molecule | 77 |
| Hs00610298_m1 | *FGF10* | Fibroblast growth factor 10 | 70 |
| Hs00232764_m1 | *FOXA2* | Forkhead box A2 | 66 |
| Hs01549772_m1 | *G6PC2* | Glucose-6-phosphatase catalytic subunit 2 | 97 |
| Hs99999905_m1 | *GAPDH* | - | 0 |
| Hs01031536_m1 | *GCG* | Glucagon | 86 |
| Hs01564555_m1 | *GCK* | Glucokinase | 72 |
| Hs00157705_m1 | *GLP1R* | Glucagon like peptide 1 receptor | 78 |
| Hs00230853_m1 | *HNF4A* | Hepatocyte nuclear factor 4 alpha | 49 |
| Hs00846499_s1 | *UCN3* | Urocortin 3 | 85 |
| Hs00355773_m1 | *INS* | Insulin | 126 |
| Hs01383002_m1 | *IRX2* | Iroquois homeobox 2 | 85 |
| Hs00158126_m1 | *ISL1* | ISL LIM homeobox 1 | 57 |
| Hs00235006_m1 | *ITGA1* | Integrin subunit alpha 1 | 87 |
| Hs01116799_m1 | *KCNK1* | Potassium two pore domain channel subfamily K member 1 | 140 |
| Hs00605529_m1 | *KCNK3* | Potassium two pore domain channel subfamily K member 3 | 134 |
| Hs00761767_s1 | *KRT19* | Keratin 19 | 116 |
| Hs04419852_s1 | *MAFA* | MAF bzip transcription factor A | 107 |
| Hs00534343_s1 | *MAFB* | MAF bzip transcription factor B | 86 |
| Hs01922995_s1 | *NEUROD1* | Neuronal differentiation 1 | 110 |
| Hs01875204_s1 | *NEUROG3* | Neurogenin 3 | 127 |
| Hs00159616_m1 | *NKX2-2* | NK2 homeobox 2 | 114 |
| Hs00232355_m1 | *NKX6-1* | NK6 homeobox 1 | 93 |
| Hs00413554_m1 | *ONECUT1* | One cut homeobox 1 | 76 |
| Hs00173014_m1 | *PAX4* | Paired box 4 | 115 |
| Hs00240871_m1 | *PAX6* | Paired box 6 | 76 |
| Hs01026107_m1 | *PCSK1* | Proprotein convertase subtilisin/kexin type 1 | 96 |
| Hs00159922_m1 | *PCSK2* | Proprotein convertase subtilisin/kexin type 2 | 76 |
| Hs00236830_m1 | *PDX1* | Pancreatic and duodenal homeobox 1 | 73 |
| Hs00426805_m1 | *GP2* | Glycoprotein 2 | 75 |
| Hs00358111_g1 | *PPY* | Pancreatic polypeptide | 68 |
| Hs01560299_m1 | *SLC16A1* | Solute carrier family 16 member 1 | 95 |
| Hs00915193_m1 | *SLC18A1* | Solute carrier family 18 member A1 | 63 |
| Hs00168966_m1 | *SLC2A4* | Solute carrier family 2 member 4 | 89 |
| Hs00545183_m1 | *SLC30A8* | Solute carrier family 30 member 8 | 73 |
| Hs00751752_s1 | *SOX17* | SRY-box 17 | 149 |
| Hs00165814_m1 | *SOX9* | SRY-box 9 | 102 |
| Hs00356144_m1 | *SST* | Somatostatin | 86 |
| Hs00300531_m1 | *SYP* | Synaptophysin | 63 |
| Hs00169095_m1 | *IAPP* | Islet amyloid polypeptide | 61 |
| Hs00371661_m1 | *TSPAN1* | Tetraspanin 1 | 87 |
| Hs04260367_gH | *POU5F1* | POU class 5 homeobox 1 | 77 |

**Table S7. Thermo Fisher TaqMan Micro Array configuration for the analysis of genes associated with pluripotency.**

| Assay ID | Gene | Gene Name(s) | Amplicon Length |
| --- | --- | --- | --- |
| Hs01053790_m1 | *ABCG2* | ATP binding cassette subfamily G member 2 (Junior blood group) | 83 |
| Hs00923299_m1 | *ACVR1B* | activin A receptor type 1B | 74 |
| Hs00609603_m1 | *ACVR2B* | activin A receptor type 2B | 101 |
| Hs01029144_m1 | *ALPL* | alkaline phosphatase | 79 |
| Hs00187842_m1 | *B2M* | beta-2-microglobulin | 64 |
| Hs00204257_m1 | *CD274* | CD274 molecule | 77 |
| Hs01023895_m1 | *CDH1* | cadherin 1 | 80 |
| Hs00172106_m1 | *CIITA* | class II | 63 |
| Hs00175480_m1 | *CTLA4* | cytotoxic T-lymphocyte associated protein 4 | 93 |
| Hs00607978_s1 | *CXCR4* | C-X-C motif chemokine receptor 4 | 153 |
| Hs99999905_m1 | *GAPDH* | - | 0 |
| Hs00999691_m1 | *FGF4* | fibroblast growth factor 4 | 130 |
| Hs00915142_m1 | *FGFR1* | fibroblast growth factor receptor 1 | 62 |
| Hs00232764_m1 | *FOXA2* | forkhead box A2 | 66 |
| Hs00231106_m1 | *FOXO1* | forkhead box O1 | 103 |
| Hs01106466_s1 | *FUT4* | fucosyltransferase 4 | 152 |
| Hs00171403_m1 | *GATA4* | GATA binding protein 4 | 68 |
| Hs00220998_m1 | *GDF3* | growth differentiation factor 3 | 65 |
| Hs01058806_g1 | *HLA-A* | major histocompatibility complex | 70 |
| Hs00818803_g1 | *HLA-B* | major histocompatibility complex | 189 |
| Hs00167041_m1 | *HNF1A* | HNF1 homeobox A | 96 |
| Hs00230853_m1 | *HNF4A* | hepatocyte nuclear factor 4 alpha | 49 |
| Hs99999909_m1 | *HPRT1* | hypoxanthine phosphoribosyltransferase 1 | 100 |
| Hs00961622_m1 | *IL10* | interleukin 10 | 74 |
| Hs00174131_m1 | *IL6* | interleukin 6 | 95 |
| Hs00235006_m1 | *ITGA1* | integrin subunit alpha 1 | 87 |
| Hs01041011_m1 | *ITGA6* | integrin subunit alpha 6 | 64 |
| Hs00174029_m1 | *KIT* | KIT proto-oncogene receptor tyrosine kinase | 64 |
| Hs00358836_m1 | *KLF4* | Kruppel like factor 4 | 110 |
| Hs00702808_s1 | *LIN28A* | lin-28 homolog A | 143 |
| Hs00153408_m1 | *MYC* | v-myc avian myelocytomatosis viral oncogene homolog | 107 |
| Hs04260366_g1 | *NANOG* | Nanog homeobox | 99 |
| Hs00240871_m1 | *PAX6* | paired box 6 | 76 |
| Hs00236830_m1 | *PDX1* | pancreatic and duodenal homeobox 1 | 73 |
| Hs01574644_m1 | *PODXL* | podocalyxin like | 82 |
| Hs00210532_m1 | *PODXL2* | podocalyxin like 2 | 73 |
| Hs04260367_gH | *POU5F1* | POU class 5 homeobox 1 | 77 |
| Hs00751752_s1 | *SOX17* | SRY-box 17 | 149 |
| Hs01053049_s1 | *SOX2* | SRY-box 2 | 91 |
| Hs00165814_m1 | *SOX9* | SRY-box 9 | 102 |
| Hs00266645_m1 | *FGF2* | fibroblast growth factor 2 | 82 |
| Hs00911929_m1 | *TBX2* | T-box 2 | 60 |
| Hs00972656_m1 | *TERT* | telomerase reverse transcriptase | 79 |
| Hs00907219_m1 | *TPBG* | trophoblast glycoprotein | 100 |
| Hs00864535_s1 | *UTF1* | undifferentiated embryonic cell transcription factor 1 | 102 |
| Hs01938187_s1 | *ZFP42* | ZFP42 zinc finger protein | 146 |
| Mr04269880_mr | *SEV* | Sendai | 59 |
| Mr04421257_mr | *SEV-KOS* | Sendai-KLF4-KOS | 80 |
