## Supplementary figures and images for "AT7867 promotes pancreatic progenitor differentiation of human iPSCs and accelerates diabetes reversal"

### Supplementary figures

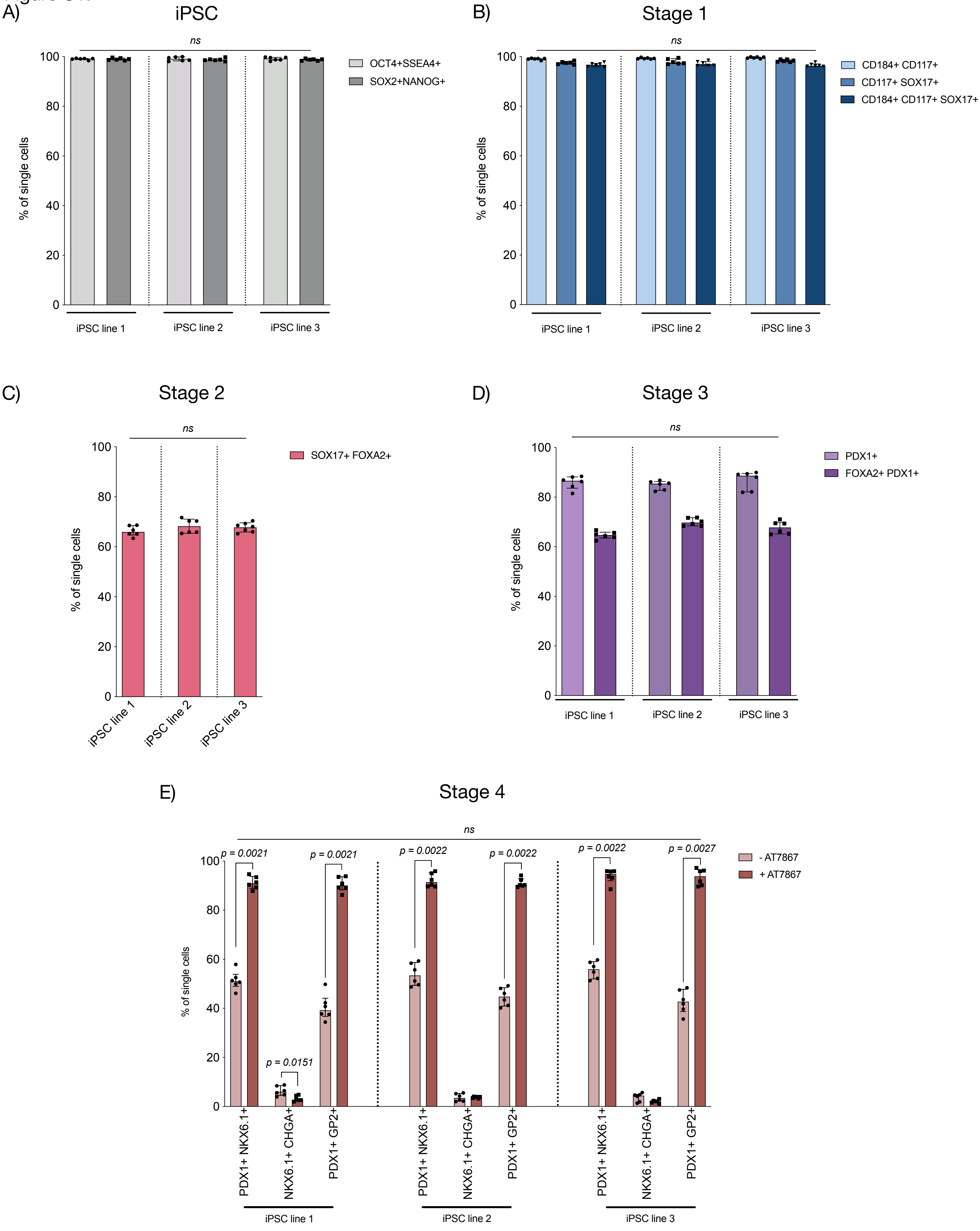

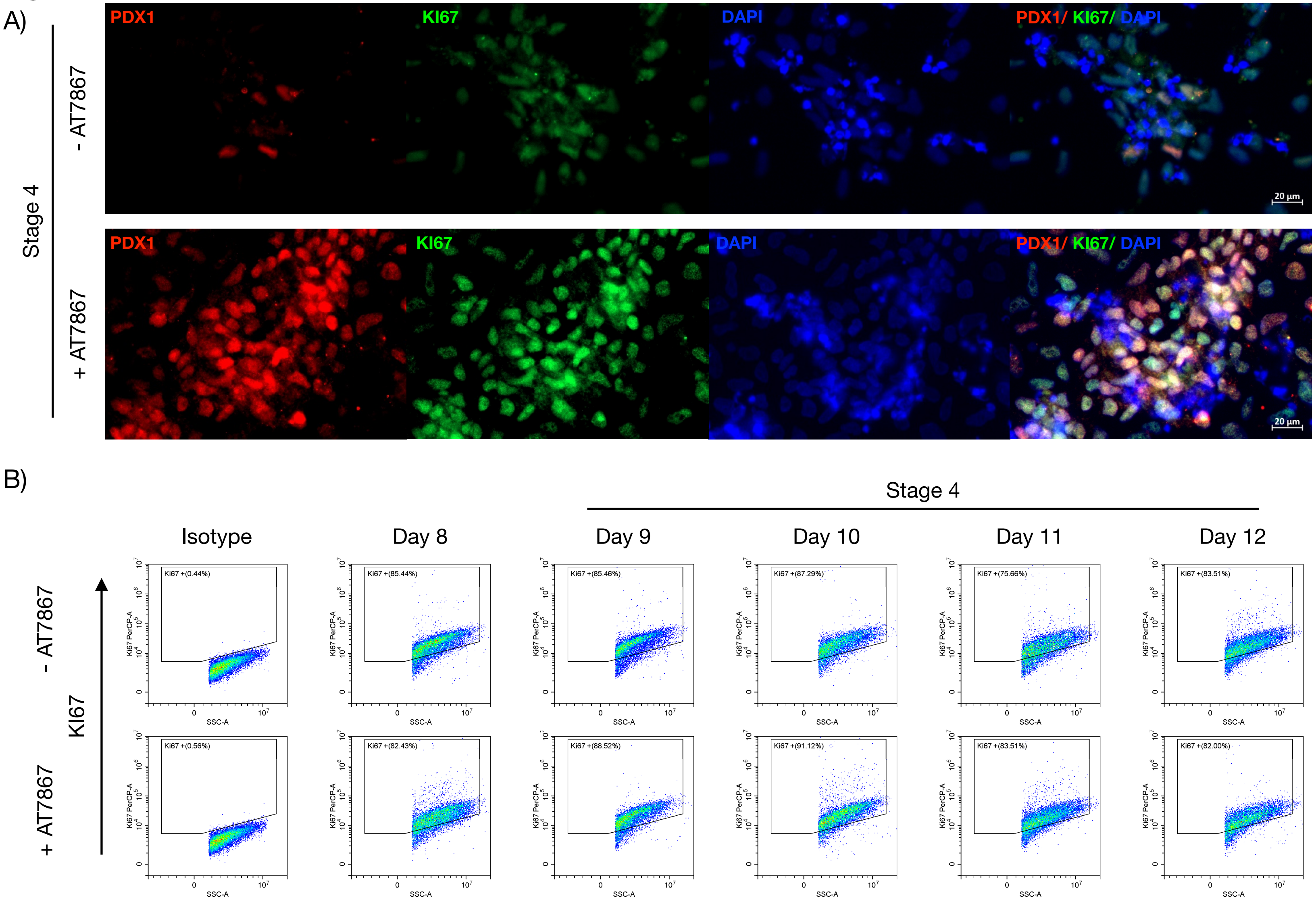

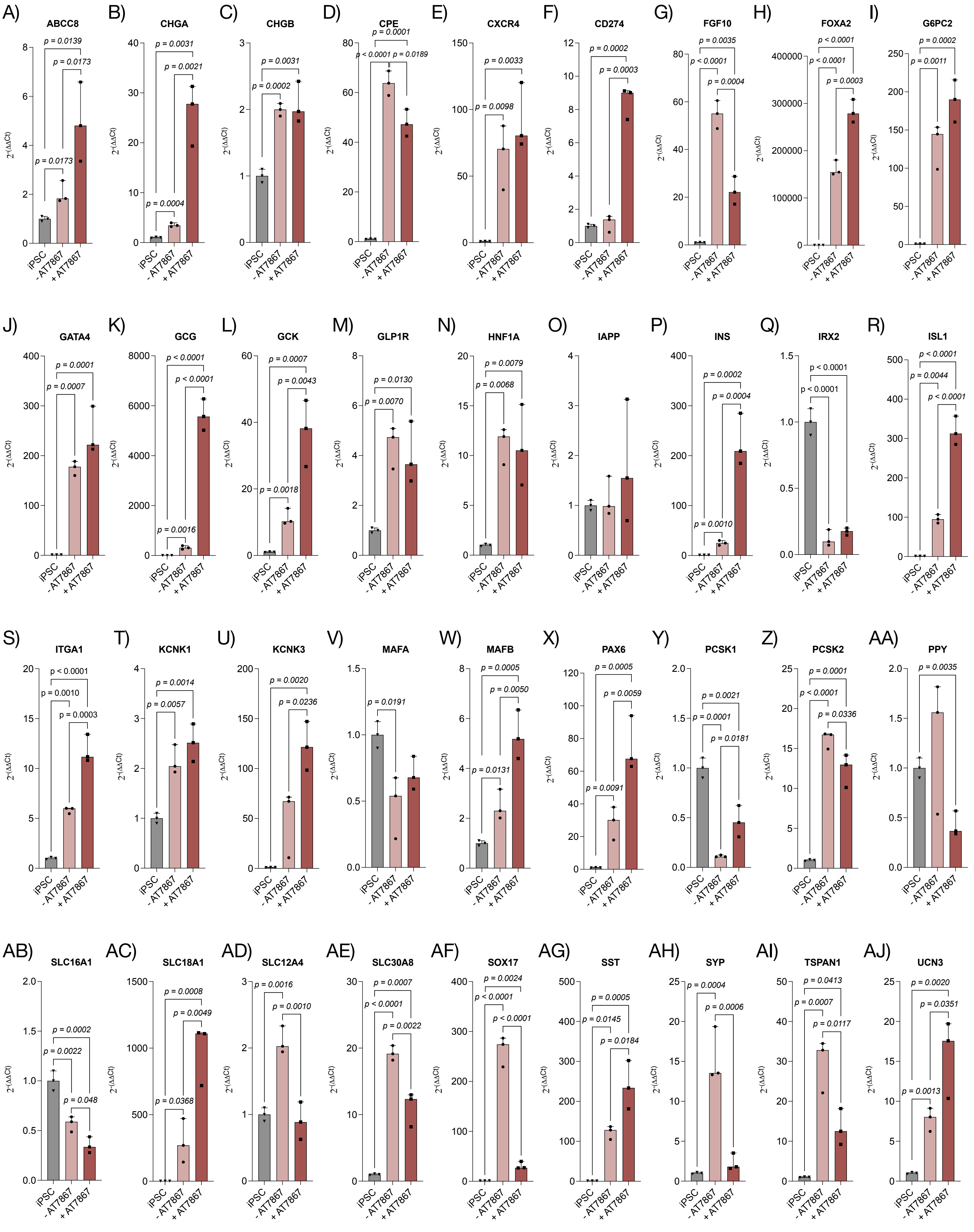

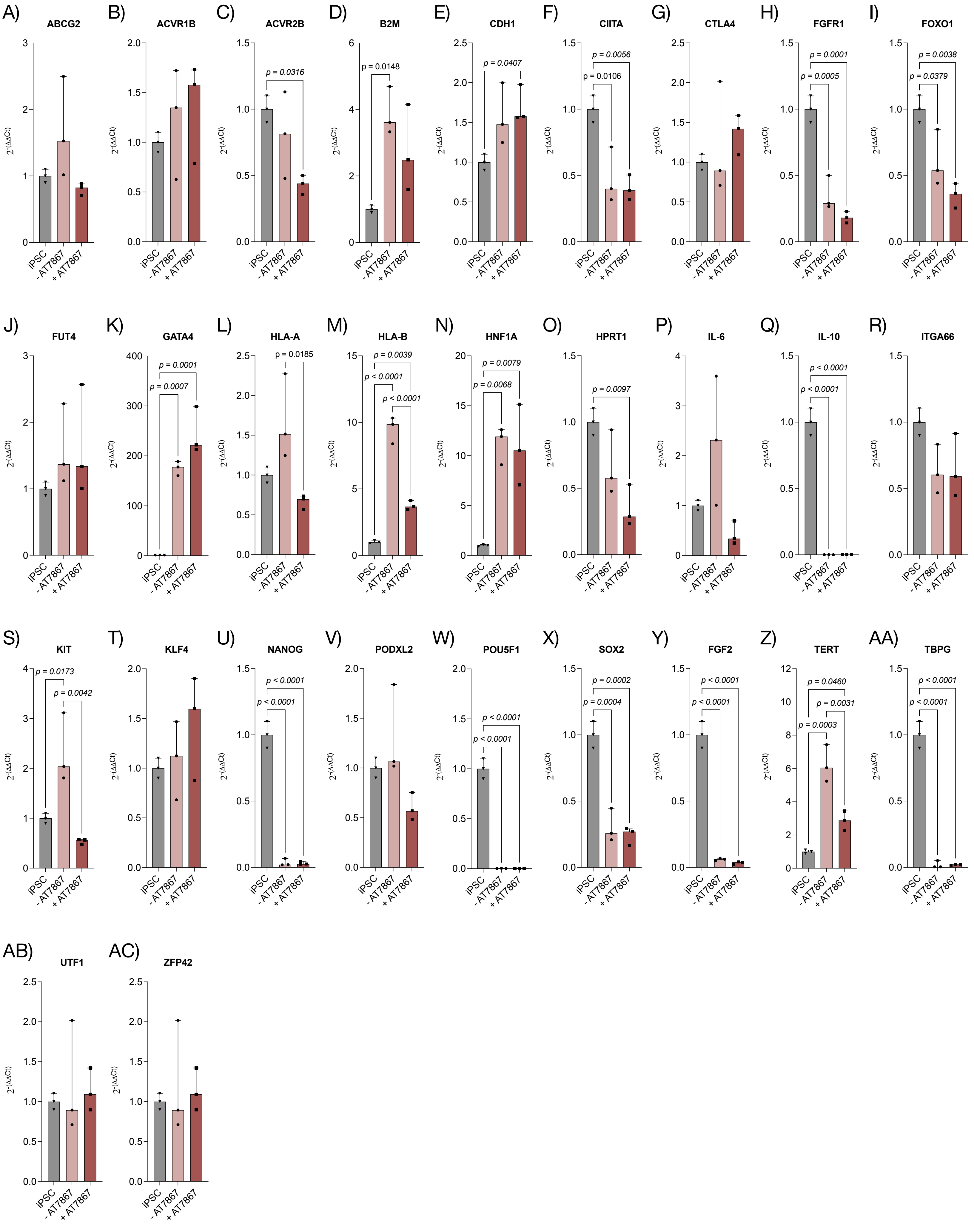

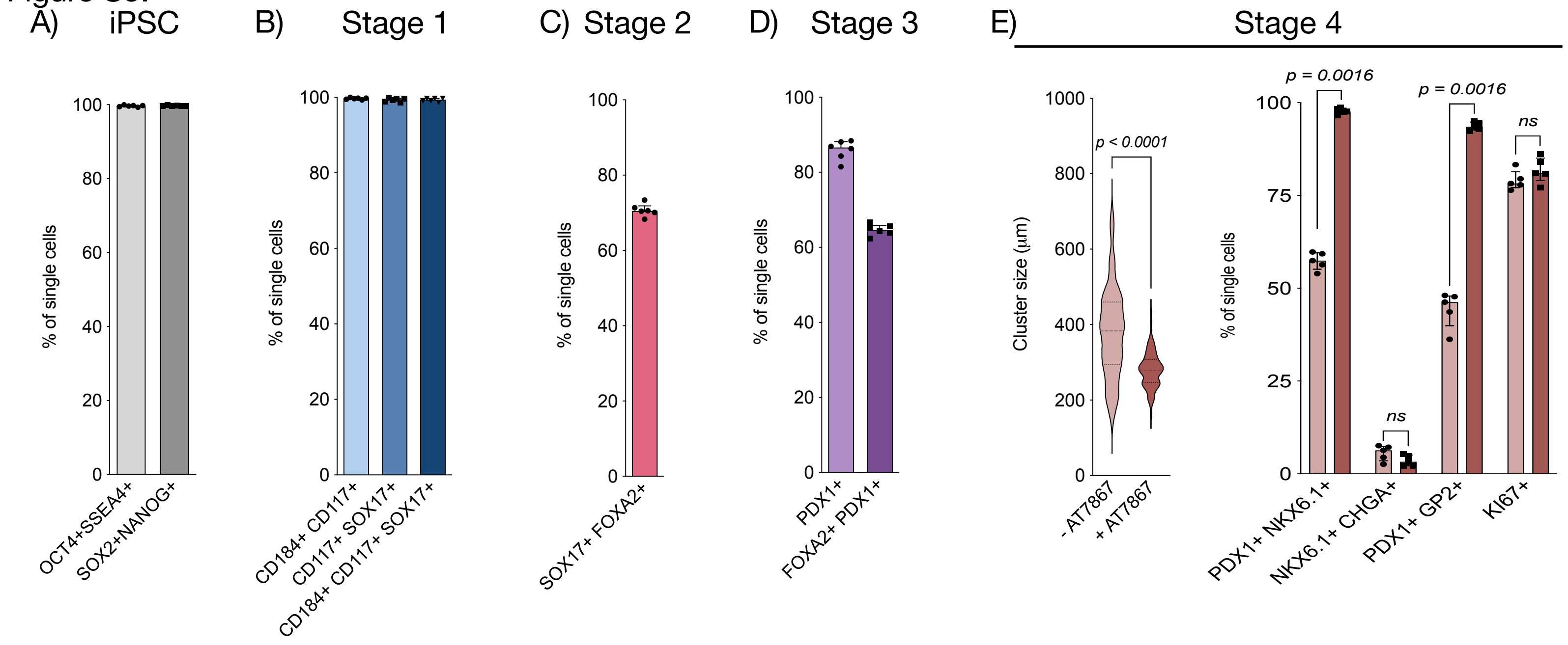
